# Validation and Long-Term Follow Up of CD33 Off-Targets Predicted In Vitro and In Silico Using Error-Corrected Sequencing in Rhesus Macaques

**DOI:** 10.1101/2020.07.05.186858

**Authors:** Aisha A. AlJanahi, Cicera R. Lazzarotto, Shirley Chen, Tae-Hoon Shin, Stefan Cordes, Isabel Jabara, Yifan Zhou, David Young, Byung-Chul Lee, Kyung-Rok Yu, Yuesheng Li, Bradley Toms, Ilker Tunc, So Gun Hong, Lauren L. Truitt, Julia Klermund, Miriam Y. Kim, Toni Cathomen, Saar Gill, Shengdar Q. Tsai, Cynthia E. Dunbar

**Author notes:** Correspondence to: Cynthia E Dunbar, Building 10-CRC, 5E-3332, NIH, 9000 Rockville Pike, Bethesda, MD 20892.

## Abstract

The programmable nuclease technology CRISPR/Cas9 has revolutionized gene editing in the last decade. Due to the risk of off-target editing, accurate and sensitive methods for off-target characterization are crucial prior to applying CRISPR/Cas9 therapeutically. Here, we utilized a rhesus macaque model to ask whether CIRCLE-Seq (CS), an *in vitro* off-target prediction method, more accurately identifies off-targets compared to *in silico* prediction (ISP) based solely on genomic sequence comparisons. We use AmpliSeq HD error-corrected sequencing to validate off-target sites predicted by CIRCLE-Seq and ISP for guide RNAs designed against *TET2* and *CD33* genes. A gRNA targeting TET2 designed using modern algorithms and predicted to have low off-target risk by both ISP and CIRCLE-Seq created no detectable mutations at off-target sites in hematopoietic cells following transplantation, even when applying highly sensitive error-corrected sequencing. In contrast, a *CD33* gRNA designed using less robust algorithms with over 10-fold more off-targets sites predicted by both ISP and CIRCLE-Seq, however there was poor correlation between the sites predicted by the two methods. When almost 500 sites identified by each method were searched for in hematopoietic cells following transplantation, 19 detectable mutations in off-target sites were detected via error-corrected sequencing. Of these 19 sites, 8 sites were predicted in the top 500 sites by both methods, 8 by CIRCLE-Seq only, and 3 by ISP only. Cells with off-target editing exhibited no expansion or abnormal behavior *in vivo* in animals followed for up to 2 years. In conclusion, neither methodology predicted all sites, and a combination of careful gRNA design, followed by screening for predicted off-target sites in target cells by multiple methods may be required for optimizing safety of clinical development.

## INTRODUCTION

Cas9 nucleases can be programed by guide RNAs (gRNA) to induce double stranded DNA breaks (DSB) in a genomic locus of interest^1–3^. The ease at which this nuclease can be programmed and delivered to cells makes it an ideal system for gene therapies^4,5^. However, Cas9 genome editors have been shown to result in unintended off-target editing^6–10^. Accurate predictive methods for CRISPR/Cas9 off-target site identification are crucial prior to applying such systems therapeutically.

*In silico* prediction algorithms (ISP) rely solely on sequence similarity between genomic loci and the gRNA. Sites similar to the guide are scored based on knowledge of gRNA and DNA binding dynamics^11,12^. Even though ISP could be customized to patient genomes if genotyping is available, it is typically performed on reference genomes and is not specific to any individual patient or animal. Cellular off-target prediction methodologies such as GUIDE-seq have been devised based on the principle of efficient integration of an end-protected DNA tag, followed by tag-specific amplification and sequencing to identify a list of off-target sites^13^. Although cellular assays are directly relevant to real off-target editing in living cells, they are challenging to perform on certain primary cell types. Hematopoietic stem cells (HSCs), for example, are very sensitive to high concentrations of editing components and to DSB. Even though HSCs can withstand the levels of editing resulting in efficient modification of the on-target site, these cells exhibit cytotoxicity precluding use of this methodology where large concentrations of DNA editing components are introduced into the cell for sensitive off-target detection.

To overcome the sensitivity limitations of cellular assays and potential lack of biologic specificity of ISP, *in vitro* off-target prediction methods were developed^14^. These methods rely on exposure of naked genomic DNA to high concentrations of Cas9 and gRNA ribonucleoprotein (RNP). *In vitro* methods bypass the cellular delivery requirement, allowing for the saturation of the reaction, which increases sensitivity and reproducibility^14^.

To date, these off-target prediction methods have been validated in cell lines or *in vivo* on only small numbers of ISP-predicted sites. Cell lines often have abnormal genomic composition and do not serve as representative models for human therapeutics. In order to better understand and prevent risks related to off-target editing prior to clinical applications of gene editing, the predictive power of these methods must be comparatively assessed in edited cells present *in vivo* in relevant animal models. Additionally, the impact of *bona fide* off-target editing should be tracked long term to ensure that there are no adverse effects caused by unintended off-target editing.

In the current study, we compare the predictive power of CIRCLE-seq^14^, a widely-used semi-quantitative *in vitro* off-target prediction method, to ISP, specifically a commonly-utilized algorithm (http://crispr.mit.edu/about) adapted to the 2015 release of the rhesus macaque genome (MMUL8.0)^15,16^. This algorithm only scores sites that have a 4 bp or less mismatches from the guide RNA. This ISP method has been used previously in our studies and was successful in predicting *bona fide* off-targets in rhesus cells^16^. We focused on the rhesus macaque autologous transplantation model for study based on similarities of rhesus and human hematopoiesis and the strong predictive value of this model for hematopoietic stem cell gene therapies^17,18^. Hematopoietic stem and progenitor cells are electroporated with RNPs and reinfused into the autologous animal following conditioning irradiation^5^ (Figure S1). We selected gRNAs targeting genes with either great biologic (TET2) or clinical (CD33) relevance. Loss-of-function mutations in TET2 have been linked to clonal hematopoiesis of aging and we have used CRISPR/Cas9 editing to create a rhesus macaque model of this clinical syndrome^19^. Creation of CD33 loss-of-function mutations in HSPC is being explored as an approach to protect normal bone marrow function from CAR-T cells targeting myeloid leukemias^5^. Following engraftment, blood and other hematopoietic tissues can then be monitored long-term for editing at on target sites and at off-target sites predicted by ISP or CIRCLE-Seq.

## RESULTS

### Identification and comparison of off-target sites predicted via *in vitro* CIRCLE-Seq or *in silico* algorithm

We selected four gRNAs currently being actively utilized in our program to edit rhesus macaque CD34+ HSPCs, specifically targeting sites in *PPP1R12C* (containing the AAVS1 “safe harbor” intronic sequence), *DNMT3A, TET2* and *CD33* genes^5,16^. The AAVS1 guide was initially reported in 2013^20^. *DNMT3* and *TET2* gRNAs were designed using the Benchling webtool for gRNA design for the rhesus macaque genome (https://www.benchling.com/crispr/). The CD33 guide was selected based on its resemblance to its human homolog with only one bp mismatch between the human and rhesus guides. The human CD33 gRNA was selected due to its efficacy and was truncated to 18bp based on reports suggesting that a truncated gRNA is more specific^5,21,22^.

We applied a previously published *in silico* ISP algorithm (http://crispr.mit.edu/about) to the 2015 release of the rhesus macaque genome (MMUL8.0) (http://www.ensembl.org, release 89) to each gRNA^15^. This algorithm includes sites that have up to 4 bp mismatches from the guide. It was successful in predicting *bona fide* off-target sites in rhesus macaque cell lines and was the most widely utilized ISP algorithm available at the time this study was initiated^16^. We performed CIRCLE-Seq for each guide on rhesus macaque blood cell DNA^14^. In brief, DNA was sheared, circularized and exposed to high concentrations of RNP complexes. Linearized molecules were then sequenced to create a list of cleavage sites, ranked by the number of reads.

For the four selected gRNAs, the overall number of CIRCLE-Seq and ISP sites correlated well between CIRCLE-Seq and ISP methodologies (Figure 1A). Of the gRNAs tested, *TET2* had the lowest number of off-target sites predicted by both methods (ISP:196, CS:36) and the *CD33* guide had the highest number (ISP:2587, CS:871). The *TET2* and *CD33* gRNAs were selected for further investigation.

**Figure 1.**
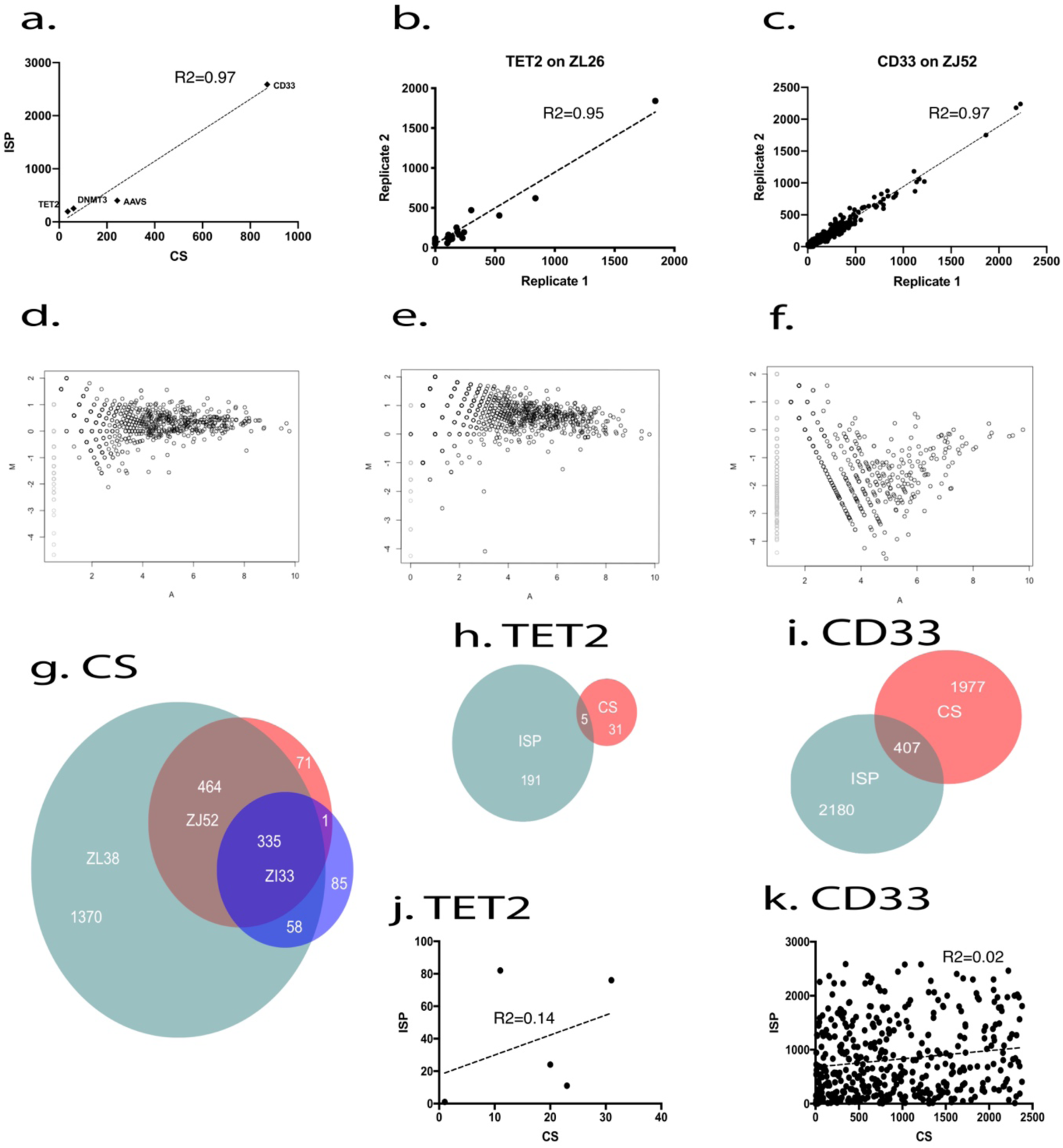
Optimizing CIRCLE-Seq. (a) Number of sites predicted for 4 different guides using CIRCLE-Seq and ISP. (b) a plot of the normalized read counts of the CIRCLE-seq detected sites for TET2 replicates. (c) A plot of the normalized read counts of the CIRCLE-seq detected sites for CD33 replicates. (d-f) Visualization of differences between the predicted site with an MA plot, where M is the log ratio scale and A is the mean average scale (average read count for each site in all 3 runs). (g) Overlap between the CIRCLE-Seq sites detected for each of our CD33 animals. (h) Overlap between TET2 sites predicted with CIRCLE-Seq on ZL26 and ISP. (i) Overlap between CD33 sites predicted with CIRCLE-Seq on 3 animals and ISP. (j,k) Spearman correlation plots of rank for the off-target sites that were predicted by both methods, CIRCLE-Seq and ISP.

To confirm the reproducibility of CIRCLE-seq, we performed technical CIRCLE-Seq replicates with *TET2* and *CD33* gRNAs on unedited DNA from animals ZL26 and ZJ52, respectively. The correlations between replicates were very high (R^2^ ≥ 0.950) (Figure 1B-C, Figure S2). Sites not identified in both replicates had very low read counts.

Next, to define the impact of genetic variation on Cas9 genome-wide activity, we performed CIRCLE-seq on genomic DNA from 3 animals using the CD33 gRNA. We normalized CIRCLE-seq read counts to the on-target site for each replicate. SNPs Read counts were again normalized for each of the three animal runs and off-target sites for all three animals were compared (Figure 1D-G). Despite differences in sequencing depth (2 replicates with a total of >16 million reads for animals ZL38 and ZJ52, and 1 replicate with a little over 4 million reads for ZL33) and efficiency of CIRCLE-Seq between runs on the DNA from the three animals, the top sites with high read counts were detected and ranked high for all three animals and pairwise comparisons showed reasonable correlations (R^2^ > 0.62) (Figure S3). In fact, the top 50 CIRCLE-Seq sites were predicted for all three animals, and of the top 200 sites, only 34 were not predicted for all three animals (Figure 1D-F). 6 of the 34 sites had unique SNPs that could explain these differences in detection. The CIRCLE-Seq sites predicted for all three animals were then combined in a CD33 master list.

We compared CIRCLE-Seq versus ISP predicted off-target sites. Only 5 of the 36 CIRCLE-Seq and 196 ISP sites for the *TET2* gRNA overlapped, and only 407 of 2384 CIRCLE-Seq and 2587 ISP sites for the *CD33* gRNA overlapped (Figure 1 H, I). Spearman correlations comparing ranking of ISP versus CIRCLE-Seq sites were very low for both *TET2* (R^2^=0.14) and *CD33* (R^2^=0.02) gRNAs (Figure 1 J, K).

### Pilot validation of top CIRCLE-Seq and ISP *CD33* gRNA off-target sites in rhesus macaque hematopoietic cells via targeted standard Illumina sequencing

We performed a pilot experiment to validate *bona fide* off-target editing in CD34+ HSPCs edited with RNPs targeting *CD33* and removed from the “infusion product” (IP) used for autologous transplantation of macaque ZJ52. Primers for the 15 top sites ranked by read count from CIRCLE-Seq performed on DNA from animal ZJ52 and the 15 top sites ranked by ISP for the *CD33* gRNA were prepared and used to amplify and then sequence these sites via standard Illumina procedures to a depth of at least 200,000 reads (Table 1). The *CD33* on-target site had 99.8% editing in the infusion product. Five of the predicted off-target amplified regions had mutated reads of greater than 1%. Three of these sites were predicted by both CIRCLE-Seq and ISP (ZJ52-CS1/ISP7, ZJ52-CS3/ISP8 and ZJ52-CS25/ISP1) and had typical insertions and deletions (indels) centered around the predicted cut site, especially in the topmost common sequencing reads (Tables S1-S3). Two sites (ZJ52-CS9/ISP1258 and ZJ52-CS13) instead had changes consisting of single or a few deleted nucleotides, often not centered at a predicted cut site, and distributed evenly across sequencing reads, suggesting a sequencing artifact rather than *bona fide* off-target editing (Tables S4, S5).

**Table 1.**
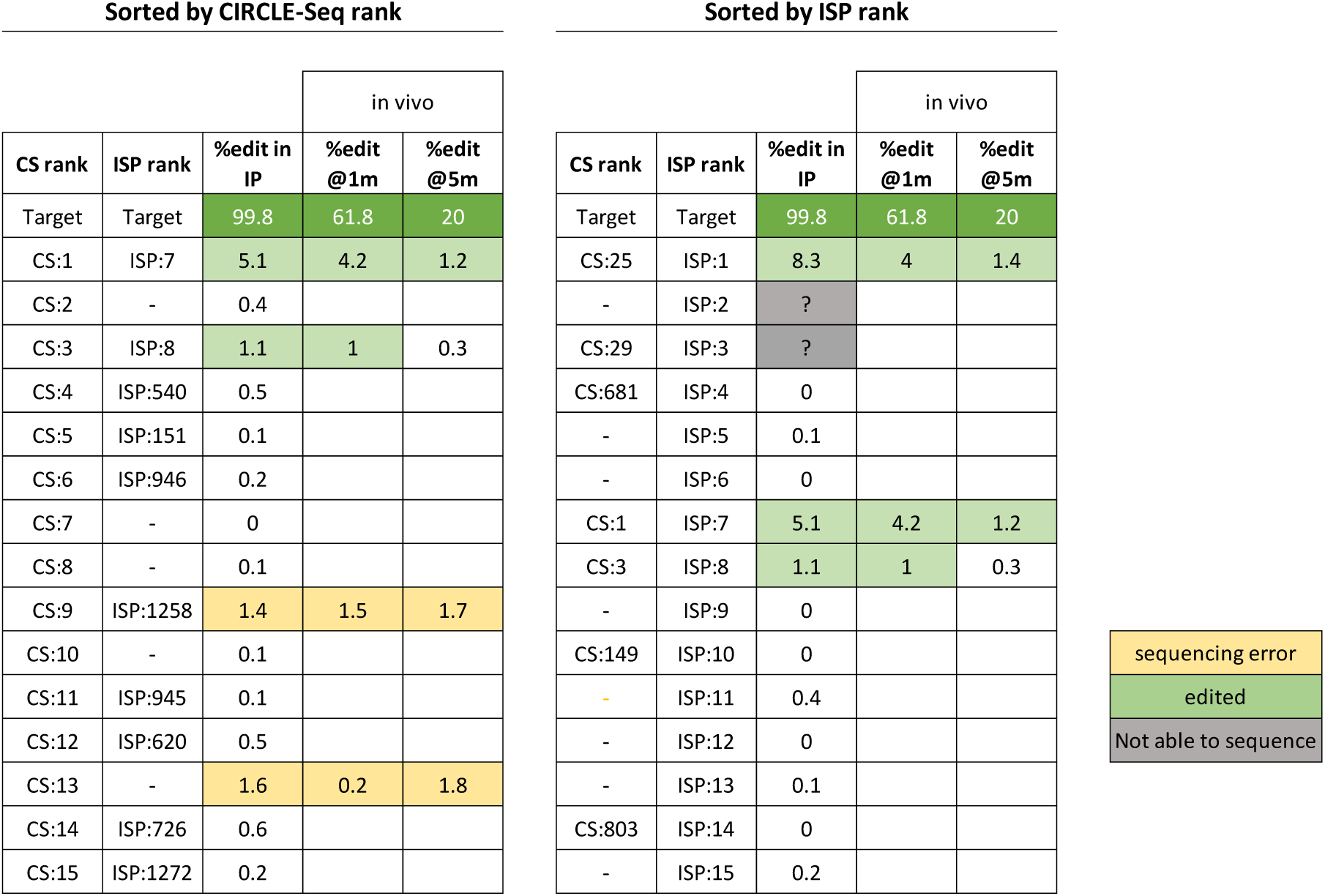
Results of sequencing the top15 CIRCLE-Seq sites predicted for ZJ52 with the CD33 guide (left) and top 15 ISP sites for the same guide (right).

Editing of the five sites identified in the CD34+ IP were next assessed in granulocytes collected from ZJ52 at one and five months following autologous transplantation. Granulocytes turn over every several days and thus reflect ongoing production from HSPCs. Sites ZJ52-CS1/ISP7, ZJ52-CS3/ISP8 and ZJ52-CS25/ISP1 with *bona fide* indels detected in the IP also had convincing indel patterns in the granulocyte samples from the transplanted macaque, with higher indel percentages at 1 month compared to 5 months, a similar pattern observed for the on target site in this animal and others transplanted in our program, reflecting more efficient editing of short-term as compared to long-term engrafting HSPCs^5,23^. Sites ZJ52-CS9/ISP1258 and ZJ52-CS13 again had mutations not typical for Cas9-mediated editing, and did not change between 1 and 5 months, further suggesting that these two sites were not actually edited but instead were sites prone to sequencing errors (Table 1).

This pilot analysis confirmed that both CIRCLE-Seq and ISP were able to predict valid off-target sites present in a clinically-relevant setting following autologous transplantation of edited HSPCs. However, standard Illumina sequencing resulted in ambiguous results for some sites due to likely sequencing artifacts and could not reliably be used for detection of edits present at allele fractions of less than 1%. Therefore, we moved to error-corrected sequencing for large scale analysis of a larger set of CIRCLE-Seq and ISP predicted sites.

### Application of error-corrected sequencing for validation of off-target sites identified by CIRCLE-Seq and ISP

Due to our observation of error rates higher than 1% with standard Illumina sequencing when searching for off-target editing at potential sites in primary target cells, we developed an error-corrected AmpliSeq-HD targeted custom panel. This approach to error-corrected sequencing facilitates sensitive mutation detection by adding a unique molecular index (UMI) tag to each amplicon. The UMI-tagged amplicons are then redundantly amplified to create “molecular families”. After sequencing, molecular families can be identified computationally by their UMI tags. Only if the majority of the members of a molecular family share the same sequence (e.g., majority have an indel or SNP in the same position), it is determined to be a true variant. Otherwise, it is almost certainly a sequencing error (Figure 2A).

**Figure 2.**
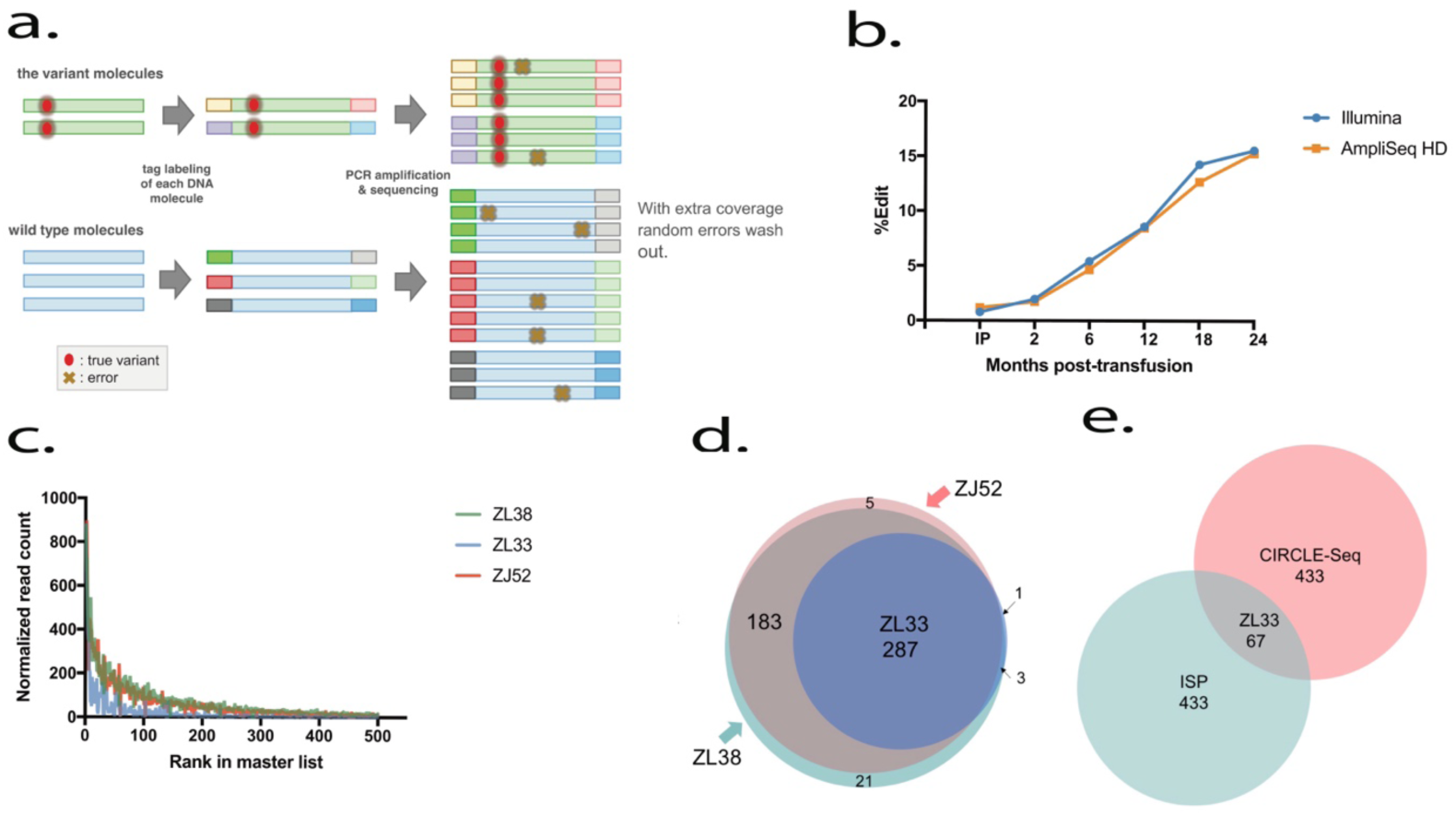
AmpliSeq HD error-corrected sequencing panels. (a) AmpliSeq HD schematic. (b) Follow up of the on-target editing in our TET2 animal ZL26 with Illumina targeted sequencing and AmpliSeq HD. (c) A plot of the read counts of the 500 selected CIRCLE-Seq sites in each animal individually. (d) The overlap of the 500 CIRCLE-Seq sites selected for AmpliSeq HD. (e) overlap of the selected 1000 sites for AmpliSeq HD in CIRCLE-Seq and ISP.

We initially created AmpliSeq HD panels for the off-target sites identified via CIRCLE-Seq and ISP for the *TET2* gRNA. The panel surveying 76 sites included all 36 *TET2* off-target sites predicted by CIRCLE-Seq and the top 40 (20%) of the sites ranked by ISP. We sampled granulocytes from animal ZL26 for 2 years post-transplantation of edited HSPC, first comparing the on-target site indel fractions obtained by AmpliSeq HD versus standard Illumina sequencing (Figure 2B). The indel fractions obtained by the two methods were nearly identical for on-target editing in the *TET2* gene at all time points (R^2^=0.99), both showing clonal expansion of HSPC with loss-of-function indels at this site. No off-target editing was detected for any CIRCLE-Seq or ISP sites using this AmpliSeq HD panel, confirming the very low off-target editing risk predicted by both CIRCLE-Seq and ISP compared to the *CD33* gRNA. AmpliSeq performed on lymphoid cells from this animal at 22 months post transplantation also demonstrated no detectable off-target editing.

We then created an AmpliSeq HD panel including the top 500 sites from the CIRCLE-Seq master list and the top 500 sites predicted by ISP for more promiscuous *CD33* gRNA. To verify whether selection of only the top 500 CIRCLE-Seq sites based on combined read rank from the master-list of CIRCLE-Seq sites predicted by runs on DNA from three animals was adequate, we plotted the read counts for a site from each individual CIRCLE-Seq run (Figure 2C). All sites with high read counts retrieved from a CIRCLE-Seq run on any animal’s DNA were ranked high on the master list. In fact, by selecting the top 500 sites, we included all CIRCLE-Seq sites that had 25 reads or more for any CIRCLE-Seq run on an individual animal’s DNA. Not surprisingly, Venn diagrams confirmed the overlap between these 500 sites in the CIRCLE-Seq runs on DNA from three animals (Figure 2D). However, only 67 sites overlapped between the top 500 CIRCLE-Seq sites and top 500 ISP sites, including the on-target site (Figure 2E2E). We were unable to design appropriate primers for 28 sites. Sites CS31, CS51 and ISP61 are the only missed sites that were in the top 100 of either method. Thus, the final AmpliSeq HD panel consisted of a total of 906 sites for *CD33* off-target analysis.

Even working with data from error-corrected sequencing, application of criteria to identify real off-target editing within the very large sequencing datasets is necessary. The error rate for AmpliSeq HD is estimated to be 5-10 fold lower than for standard Illumina sequencing, but it is not zero. We applied the following criteria to score a site as truly edited in a cell sample:

1. Only INDELS were considered as edited, given that non-homologous end joining (NHEJ) very rarely results in SNPs, and most sequencing errors mimic SNPs ^24,25^.
2. Only INDELS within a 20bp window 5’ of a PAM sequence were considered edited.
3. Baseline non-edited cells analyzed using the same AmpliSeq HD panel could not contain the edited site.
4. INDEL rate at the site is higher than 0.05%.
5. Meets any 1 of the following criteria:
  a. Molecular coverage more than 1 (has at least 2 UMIs).
  b. Present in multiple animals or multiple time points.
  c. Multiple edit types at one site.
  d. Non-repetitive INDEL > 5 bp in length.

These criteria were designed to be conservative in order to avoid false positives. However, as this is the most sensitive sequencing method available, we have no means of confirming that all sites that meet our criteria were truly edited. Nonetheless, we assume that any false positives or negatives will not be biased towards CIRCLE-Seq or ISP.

We sequenced granulocyte samples over time from four macaques transplanted with HSPCs edited with the *CD33* gRNA. Details of HSPC editing, transplantation and follow-up have been previously published for two of the animals, and Table S6 summarizes relevant information for the entire cohort^5^. By applying these criteria for the AmpliSeq HD panel data from the granulocyte samples, we detected editing at 17 predicted off-target sites. Of note, all 17 of these sites showed the same trend as the on-target site in terms of editing levels, with higher contributions at earlier time points following engraftment, then dropping to stable lower levels as more easily edited short-term progenitors exhaust and are replaced by contributions from more difficult to edit long-term HSCs (Figure 3).

**Figure 3.**
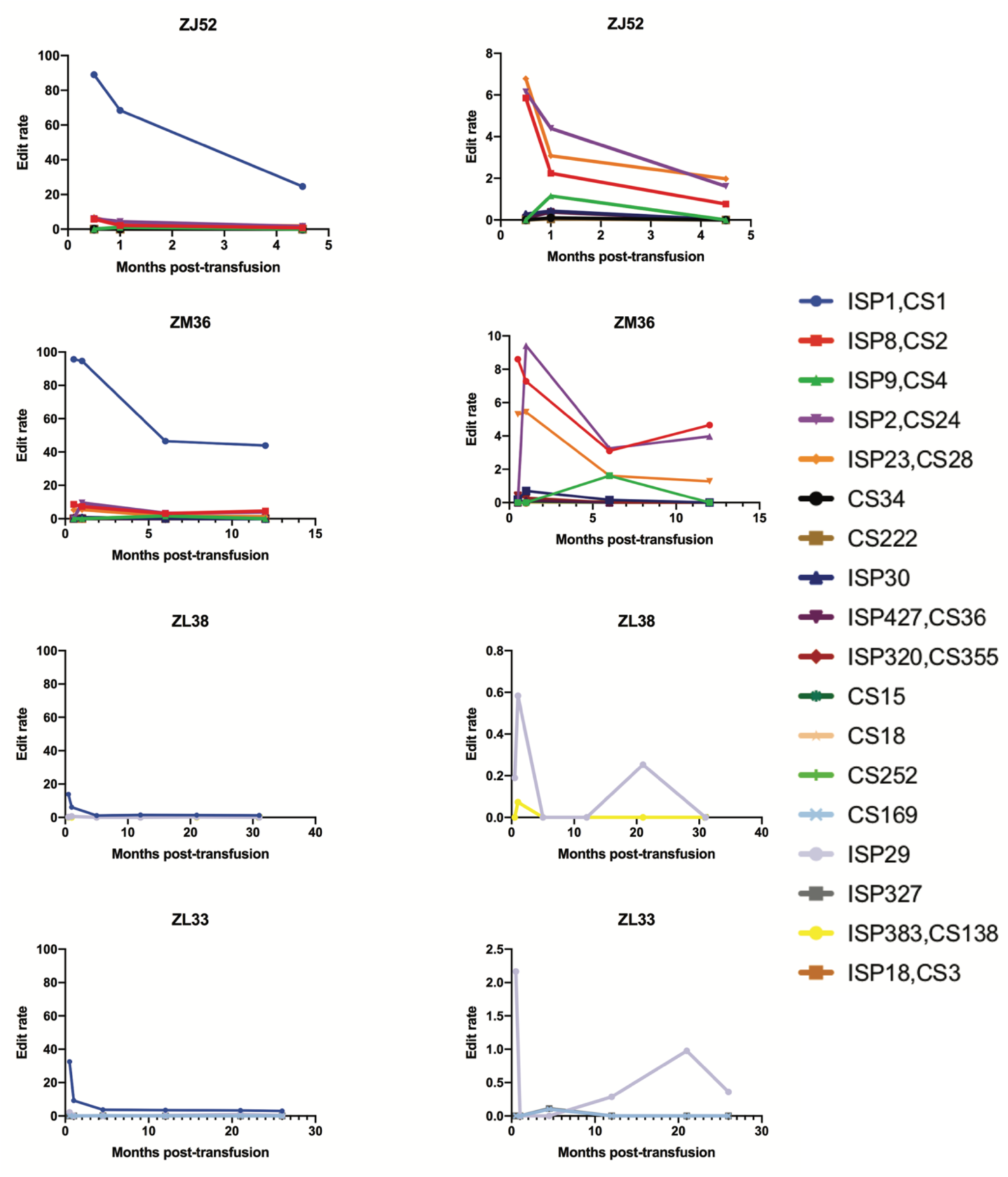
CD33 off-target edits in granulocytes over-time. The left graph for each animal shows the edited off-target sites in relation to the on-target ISP1,CS1 (in blue). The Y axis is set to a 100%. The right graph for each animal only shows the off-target sites.

In order to ensure that off-target edits potentially perturbing hematopoiesis were not being missed by focusing solely on granulocytes, we also analyzed T, B and NK cell samples from the same 4 edited animals (Table 2). We identified editing at two additional off-target sites, CS103, present at low percentages (≤ 0.15%) in T cells of animals ZL33 and ZL38, and CS391 also present at a low percentage (0.23%) in NK cells of ZJ52, thus increasing the total number of off-target sites identified as present in the animals to 19. Editing percentages in each bp of these 19 sites are shown in figure S4. About half of the off-targets sites (9/19) were detected in more than one lineage, with those off-target sites contributing at the highest levels in any lineage more likely to be found in multiple lineages, suggesting that sampling and detection limits rather than lineage-bias resulting from off-target editing accounted for these results.

**Table 2.**
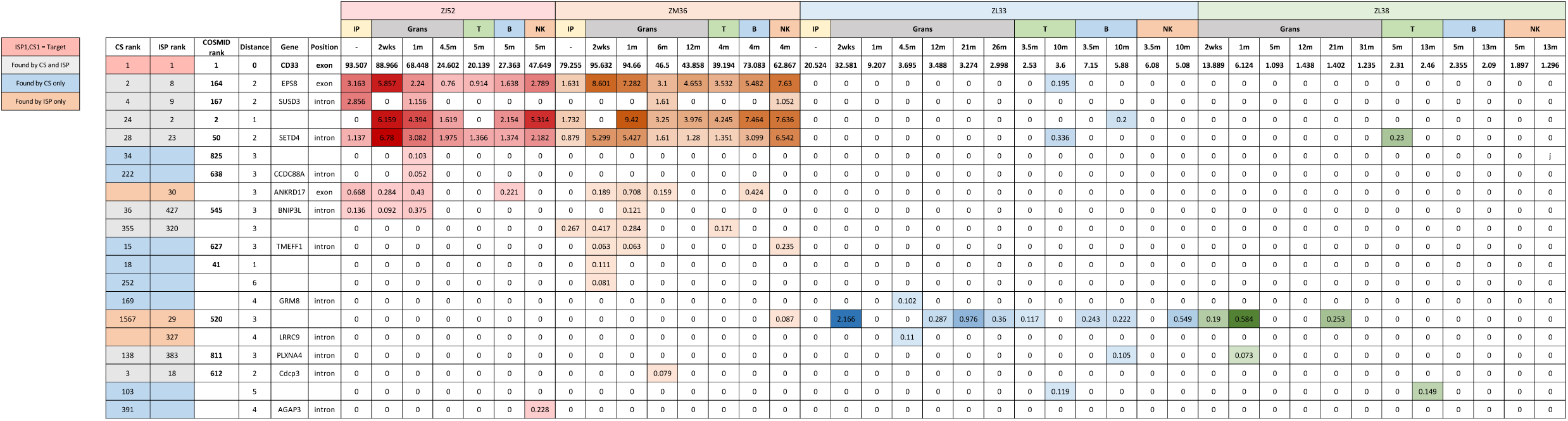
Table summary of the sequencing results of the 19 off-target sites obtained from sequencing granulocytes and T, B and NK cells in all 4 CD33 animals. ZJ52 sites are in red, ZM36 in orange, ZL33 in blue and ZL38 in green.

Many (8) of these 19 *bona fide* off-targets sites found in cells of animals following transplantation were ranked in the top 500 by both CIRCLE-Seq and ISP, despite the overall low overlap between CIRCLE-Seq and ISP predicted sites. Of the remaining sites, 8 were predicted in the top 500 by CIRCLE-Seq only, while 3 were predicted in the top 500 by ISP only. One of the ISP-only sites was predicted by CIRCLE-Seq but ranked 1567^th^ and therefore not scored as a CIRCLE-Seq top 500 site (Figure 4A). We looked at the Levenshtein Distance, which is the number of nucleotides that differ between the gRNA and the off-target site, to see if it provides any explanation as to why the 8 CS-only sites were not predicted by the ISP algorithm. 2 sites were excluded by the ISP algorithm due to having more than 4 bp mismatches to the gRNA. Additionally, 3 sites were excluded by ISP due to gaps between the DNA and gRNA, which the ISP algorithm used does not consider (Figure 4B, C). To compare our results to a more recent ISP algorithm that take gaps into account, we ran COSMID, a webtool algorithm designed to predict off-targets^26^. We found that COSMID predicted a total of 1119 off target sites for the CD33 gRNA and was able to predict 12 of the 19 *bona fide* off-target sites, however, many of them were ranked very low (Table 2). In conclusion, real off-target sites detected *in vivo* following transplantation were missed by both ISP and CIRCLE-Seq.

**Figure 4.**
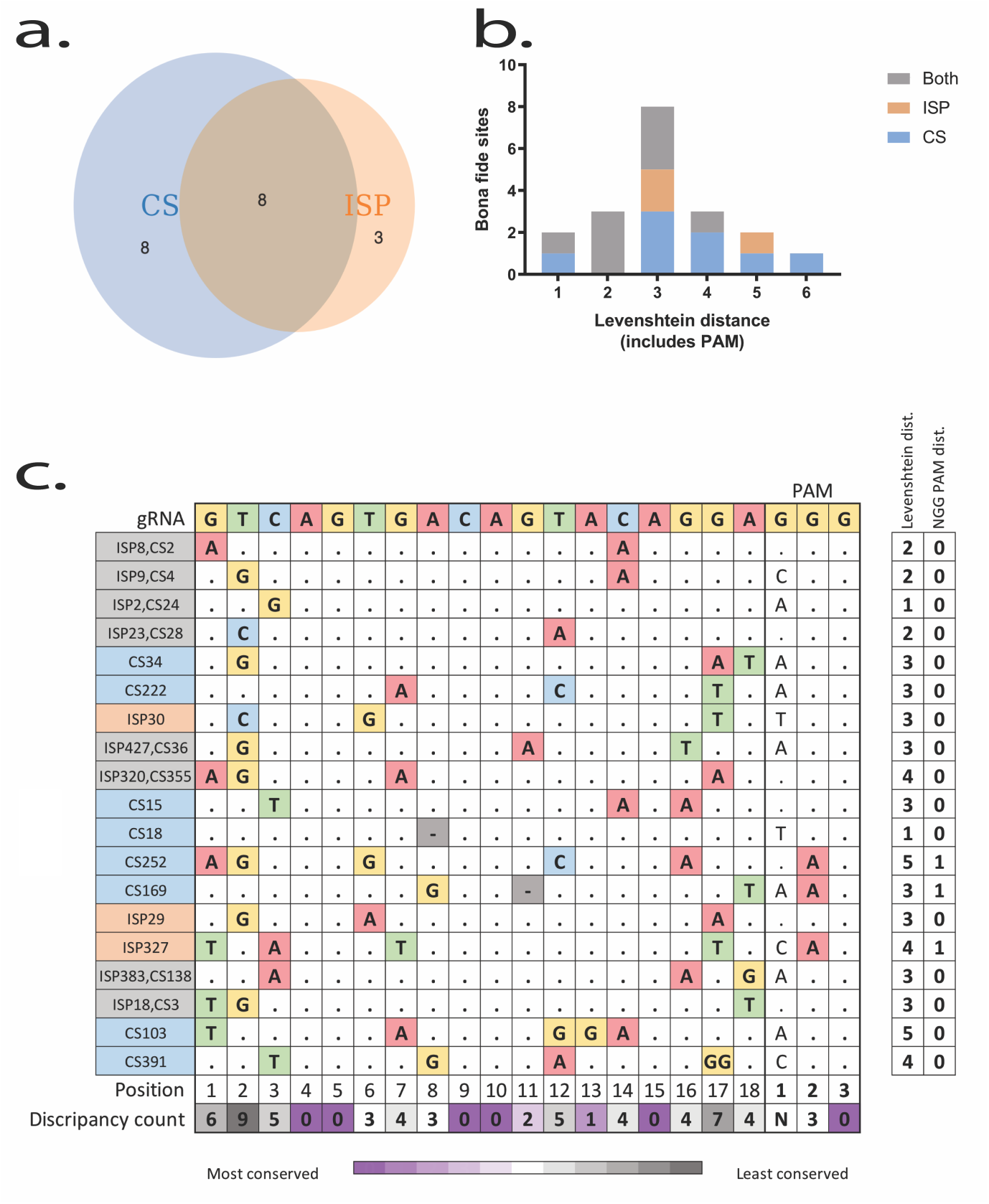
Off-target predictive power of CIRCLE-Seq vs ISP. (a) the distribution of the *bona fide* sites across the two prediction methods. (b) The Levenshtein distance between the *bona fide* off-target sites and the gRNA. (c) The types of each discrepancy from the gRNA in the *bona fide* off-target sites.

### Investigating the effects of chromatin accessibility

To understand the effect of chromatin accessibility on the presence or absence of predicted off-targets in primary edited hematopoietic cells, we compared the chromatin accessibility as assessed by available “assay for transposase-accessible chromatin” (ATAC-Seq) data derived from human HSPCs for the 1000 CD33 gRNA predicted off-target sites included on the AmpliSeq HD panel versus the 19 sites actually detected in primary hematopoietic cell samples from the macaques^27,28^. ATAC-seq data from human CD34+ cells was utilized (GEO: GSE96772) due to lack of any available data for rhesus macaque HSPCs. CIRCLE-Seq is performed on naked DNA and would thus not be predicted to be sensitive to chromatin structure, and ISP algorithms to date do not take into account any chromatin features. As described previously, the ATAC-seq peaks from human genomic coordinates (hg19) were mapped to macaque genomic coordinates (rheMac8) and predicted off-target sites were mapped onto the ATAC-seq features^29^. 51 of the set of 1000 predicted off-target sites fell within predicted open chromatin regions versus 4 of the 19 sites detected as edited in macaque samples. By chi-squared testing, there was a relationship between the presence of open chromatin and a higher likelihood of actual off-target editing (chi-square statistic= 7.283, p-value= .007 – without Yates correction). While a larger analysis of multiple gRNAs with additional statistical power will be required to strengthen claims for a relationship, these results support a connection between editing and chromatin accessibility, supporting findings in our recent paper showing the same relationship for efficiency of editing on-target sites.

## DISCUSSION

It has been challenging to design clinically-relevant approaches for the prediction and detection of off-target mutations following CRISPR/Cas gene editing. In both mice and large animals undergoing *in vivo* delivery of editing machinery to target muscle, off-target editing has not been detected, however, insensitive detection methods such as T7E1 PCR assay or non-error corrected sequencing were utilized in these earlier studies^3,30–35^. Off-target editing is potentially less challenging to detect in approaches editing embryos, given that the small number of target cells initially edited then expand to create an entire organism. Several articles reported minimal to no off-target editing in these models^7,9,36–40^, while others found high level off-target editing^41,42^. These studies, however, only screened a maximum of 10 off-target sites predicted solely using *in silico* approaches^31–35,37,38^. Since gRNAs are designed using *in silico* algorithms, predicting off-targets using the same or similar algorithms can be insensitive to sites not considered as relevant by these approaches. Additionally, none of these studies applied error-corrected sequencing, and thus could not detect off-target editing accurately at levels of less than 2-3%. Sites predicted by CIRCLE-Seq have been detected *in vivo* in mice, following editing of a locus in the liver, however, the predictive value of CIRCLE-Seq was not compared to *in silico* approaches in this prior report^3^.

Our study demonstrated that gRNA design plays a major role in determining off-target effects, whether predicted by ISP or CIRCLE-Seq. A gRNA targeting TET2 designed using modern algorithms and predicted to have low off-target risk by both ISP and CIRCLE-Seq created no detectable mutations at off-target sites in hematopoietic cells following transplantation, even when applying highly sensitive error-corrected sequencing. In contrast, a *CD33* gRNA designed using less robust algorithms with over 10-fold more off-targets sites predicted by both ISP and CIRCLE-Seq resulted in detectable off-target mutations in hematopoietic cells *in vivo* over time and in multiple lineages post-transplantation. Of note, even truncated guides, initially developed as more specific, also resulted in significant off-target editing^43,44^. We confirm that mismatches at the 5’ end of the guide were tolerated by Cas9, even when using a truncated guide (Figure 4C) ^45,46^. In addition, we utilized RNP delivery which is predicted to minimize off-target effects due to a short intracellular half-life of the editing machinery^4,30,43,47,48^. We were surprised to see a potentially different spectrum of valid off-target sites when utilizing standard *in vitro* transcribed gRNAs (animals ZL33 and ZL38) versus chemically-modified gRNAs (animals ZJ52 and ZM36). This difference could not be explained solely by the efficiency of editing. To our knowledge, similar findings have not been reported, however a published study did find a difference in the off-targets for guides transcribed with a U6 promoter as opposed to T7 promoter^12^.

Current ISP algorithms work by searching the entirety of the reference genome for sites that are similar to the gRNA, then scoring the similarity of these sites. Often, ISP is limited to sites that have 4 bp differences from the gRNA, otherwise thousands or tens of thousands of off-target sites would be predicted for every gRNA^44^. However, Cas9 has been shown to tolerate more than 4 bp mismatches and bulges between the off-target site and the gRNA^8,46,49^. Moreover, individual-specific SNP’s are not taken into account using ISP, even though SNPs have been shown to impact off-target editing^14,50^. Additionally, while ISP uses the short gRNA as query and searches for sites with mismatches to the 20 bp sequence, *in vitro* approaches have the advantage of more specific alignment due to the larger query sequence (∼300 bp for CS) without the need to allow mismatches^14^. We speculate that the few *bona fide* sites predicted by ISP but not by CIRCLE-Seq were missed either because the sites are in regions that are difficult to amplify during library preparation or difficult to sequence on the Illumina platform.

Although error-corrected sequencing is the most sensitive sequencing method available, allowing up to a 10-fold increase in sensitivity compared to standard targeted Illumina sequencing, it still has several limitations. Error-corrected sequencing is limited by the PCR error rate. Specifically, PCR polymerases are known to have problem areas and can make errors in the same position repeatedly, mimicking a mutation ^51–53^. This can result in false positives, as seen for some putative sites that were not consistent with editing. Additionally, AmpliSeq HD has only been optimized for 20 ng of DNA, which amounts to ∼3,000 cells (∼6,000 rhesus genome copies). This limits detection of low-frequency off-target sites. ISP29 in both ZL33 and ZL38 is an example of this limitation. A mutation in ISP29 is detectable in low frequencies at early time points, then disappears in the middle time points, only to appear later for both animals. We suspect that this phenomenon is due to sampling limitations that resulted in false negatives at some time points.

The major concern with off-target editing is the risk of clonal expansion and eventual neoplasia due to loss of function or change in function of a tumor repressor or a growth-control gene. Thus the impact of even rare off-target sites should be assessed prior to clinical applications^44,54^. Our AmpliSeq HD was optimized for only 20 ng of DNA (∼6,000 rhesus genome copies), which could lead to an underestimation of rare *bona fide* off-targets due to insufficient sampling. However, we believe that if clonal expansion due to a rare off-target has occurred, we would have observed it at later time points given that each macaque was transplanted with hundreds of thousands to millions of edited HSPCs. Multiple replicate AmpliSeq runs on individual samples could increase sensitivity, or the process could be optimized for higher amounts of input DNA.

Based on our findings, in HSPC engrafted in rhesus macaques, a model with high relevance to ongoing and future clinical development of CRISPR therapeutics, both CIRCLE-Seq and ISP were able to predict valid *in vivo* off-target sites, however both methods missed sites, and rankings of sites via the two approaches did not correlate well, and for either approach, ranking did not strongly predict the likelihood of the site being detected *in vivo*. While most *bone fide* CIRCLE-Seq sites could be predicted by a more liberal ISP, allowing additional mismatches, such an ISP would expand the list of potential off-target sites enormously, to levels impractical to screen for validity even with large targeted panels. Therefore, there is value in using both CIRCLE-Seq and ISP to predict off-targets and ensure that sites in cancer-linked genes are not predicted by either method. Initial selection of guide RNAs could be based on careful analysis of results from both ISP and CIRCLE-Seq, avoiding guides with worrisome sites predicted by either methodology. Large panel-based error-corrected sequences can be used to screen for *bona fide* off targets in relevant primary cells prior to choice of specific gRNAs for clinical applications.

## Materials and Methods

### Guide design

The CD33 guide was selected based on its resemblance to its human homolog with only one bp mismatch between the human and rhesus guides and its efficiency has been previously reported. It was truncated to 18bp based on reports suggesting that a truncated gRNA is more specific ^5,21,22^. The TET2 guide was designed using Benchling webtool. (https://www.benchling.com/crispr/). All guide sequences are in supplementary table S6.

### *In silico* prediction

*In silico* off-target prediction was performed with an in-house Python script modifying the previously-published and recently updated (http://crispr.mit.edu/about) algorithm for the rhesus macaque (Macaca mulatta) genome, utilizing the unmasked reference genome from Ensembl (http://www.ensembl.org, release 89) ^15,16^. This algorithm calculates a score for each potential off-target each site in the reference genome that has 4 or less bp mismatches to the gRNA.

### CIRCLE-Seq

Genomic DNA was extracted with the Gentra Puregene Kit (QIAGEN) according to manufacturer’s instructions. CIRCLE-Seq was performed as previously described ^14^. In short, genomic rhesus macaque DNA was sheared to an average of 300 bp using a Covaris S2 instrument. The sheared molecules were repaired, A-tailed and ligated to an adaptor in the form of a uracil-containing stem-loop using the KAPA HTP Library Preparation Kit PCR Free (KAPA Biosystems). Lambda exonuclease (New England Biolabs) and E. coli exonuclease I (New England Biolabs) was used to digest all molecules with free ends. The molecules were then treated with a USER-enzyme (New England Biolabs) and a T4 polynucleotide kinase (New England Biolabs). The molecules were then circularized with T4 DNA ligase (New England Biolabs). Non-circular molecules were eliminated with Plasmid-Safe ATP-dependent DNase (Epicenter). *In vitro* cleavage was performed in a 50 ul reaction consisting of 125 ng of the remaining circularized DNA molecules, 90 nM of SpCas9 protein (New England Biolabs), Cas9 nuclease buffer (New England Biolabs) and 90 nM of gRNA (either *in vitro* transcribed using the GeneArt kit (Thermofisher) or chemically modified synthetic gRNAs (Synthego). The cleaved molecules were then A-tailed and ligated with a hair-pin adaptor (New England Biolabs), treated with a USER enzyme and then amplified via PCR using universal primers Next Multiplex Oligos for Illumina (New England Biolabs) and Kapa HiFi Polymerase (KAPA Biosystems). Libraries were sequenced on an Illumina MiSeq with 150 bp paired-end reads.

### CIRCLE-Seq data analysis and normalization

Raw data was analyzed using open-source CIRCLE-seq analytic software (https://github.com/tsailabSJ/circleseq). For normalization between replicates the “identified_matched.txt” CIRCLE-Seq output files for each technical replicate were utilized and the number of reads for the on-target site with least reads comparing all replicates was used as readRef#. The multiplication factor for each technical replicate was calculated as: multiplication factor = readRef#/on target read count for that replicate. Each read count for each off-target for the replicate was then normalized as read count X multiplication factor. A python script (https://github.com/aljanahiaa/off-targets/blob/master/replicateCombiner.py) was used to combine the normalized replicates to create a combined technical replicate file. The same python script also calculated the Levenshtein distance between the guide and the predicted off-target. If the site has a different distance and/or coordinates, depending on whether a gap was allowed or not, the python script will use the distance and coordinates that allow for a gap.

In order to normalize read counts from CIRCLE-Seq between individual animals “identified_matched.txt” or combined technical replicate files for each animal were used, depending on whether technical replicates were performed for each animal or not. Similar to replicate normalization, a multiplication factor was calculated based on the smallest read number for an on-target site between animals and applied to all the read counts from that run of CIRCLE-Seq in order to normalize read counts between animals. Then, all normalized files were combined with a python script (https://github.com/aljanahiaa/off-targets/blob/master/masterlistCreator.py) and the average read count per site was calculated as such: average read count = sum of reads for site/number of animals in which this site was predicted. Calculating the average this way insured that if there are sites specific to one animal due to unique SNPs, then that site would not be penalized for not being predicted in other animals. The average read count was used to rank the sites in the master list of off-target sites for that gRNA.

### Autologous transplantation of rhesus macaques with CRISPR/Cas edited hematopoietic stem and progenitor cells

Autologous transplantation of rhesus macaques with CRISPR/Cas9 edited CD34+-enriched mobilized peripheral blood HSPCs was performed as previously described under protocols approved by the NHLBI Animal Care and Use Committee^5,55^. On Day −2, CD34+ cells were purified from the apheresis PBMC collection via immunoabsorption and cultured overnight in at 37°C in X-VIVOTM 10 (Lonza) supplemented with 1% HSA (Baxter) and cytokines (SCF 100ng/mL, FLT3L 100ng/mL and TPO 100ng/mL; all from PeproTech)^55^. The next day (day −1), RNA-protein complexes (RNPs) were prepared by mixing Cas9 protein 30-60 ug (PNA Bio) and 15-30 ug of gRNA per aliquot followed by incubation of 10 minutes at room temperature. Target CD34+ cells were removed from culture, washed with PBS, and then resuspended in aliquots of 3-5×10^6^ CD34+ cells in a total volume of 750ul of Opti-MEM (Thermo Fisher Scientific). An aliquot of RNPs were added to the cell suspension and electroporated using the BTX ECM 830 Square Wave Electroporation System (Harvard Apparatus) with a single pulse of 400V for 5msec. The electroporated cells were pooled and incubated at 32°C overnight in X-VIVOTM 10, 1% HSA and SCF, FLT3L and TPO at the concentrations given above. The autologous macaque underwent total body irradiation with 400-500 cGy/day on days −1 and 0. Several hours following TBI on day 0, the edited CD34+ cells were infused intravenously back into the animal. More information on the transplantation and editing parameters for each animal is given in Supplemental Table S1.

### Collection and purification of hematopoietic cells post-transplantation

Peripheral blood samples are layered onto Lymphocyte Separation Medium (MP Biomedicals) to separate mononuclear cells (MNCs) and granulocytes. Red blood cells in each fraction were lysed with ACK lysis buffer (Quality Biological). The MNCs were stained for flow cytometric using antibodies described in supplemental table S7. T cells, B cells and NK cells were isolated via fluorescence-activated cell sorting on a BD FACSAria II instrument.

### AmpliSeq HD

The custom primer panels for the AmpliSeq HD sites were designed by the White Glove team at Thermofisher using the rheMac8 reference genome (https://www.ncbi.nlm.nih.gov/assembly/GCF_000772875.2/). Amplicons were 70-225bp for *TET2* and 59-161bp for *CD33*. The libraries were prepared using the Ion AmpliSeq™ HD Library Kit (Thermofisher) according to manufacturer’s instructions using the custom primer panels for the first PCR and the Ion AmpliSeq™ HD Dual Barcode Kit (Thermofisher) for the second PCR. Each 3 samples for *TET2* and 2 samples for *CD33* were pooled together and templated on the IonChef instrument, then sequenced on an IonTorrent S5 instrument on Ion 530 and Ion 550 chips for TET2 and CD33, respectively. The S5 Torrent Server was used to analyze the samples. Torrent Variant Caller 5.12 plug in was used with custom parameters made available at (https://github.com/aljanahiaa/off-targets/blob/master/customVariantCallerParameters.json) as a JavaScript Object Notation file. The variant caller Excel output files were downloaded and inspected manually for edits that met criteria for valid indels.

### Targeted Illumina sequencing

Targeted sequencing for specific off-target or on-target sites was performed via a standard 2-step PCR using gene-specific primers with adaptors in the first round of PCR amplification and NEBNext^®^ Multiplex Oligos for Illumina^®^ (Dual Index Primers Set 1) (New England Biolabs) for the second round of PCR amplification. Gene specific primers were designed using Primer3Plus (http://www.bioinformatics.nl/cgi-bin/primer3plus/primer3plusHelp.cgi). Most amplicons were around 250 bp in length. Forward and reverse adaptors for the NEBNext^®^ Multiplex were added 5’ of the gene specific primers. The adaptor sequences are provided in supplementary table S8. For the first round, 12.5 ul of KAPA HiFi HotStart ReadyMix polymerase (KAPA Biosystems) and 0.75 ul of each 10uM primer was added to 20 ng of cellular DNA. For the second round of PCR, a unique combination of 1.5 ul of each i5 and i7 primers from the NEBNext kit were added plus an additional 22 ul of Kapa polymerase were added to the same tube. Cycling conditions for both PCR rounds were: 3 minutes at 95° degrees, 20 cycles of [98° for 20s, 62° for 15s, 72° for 30s] followed by 1 minute at 72°. Sequencing was performed with an Illumina MiSeq with 150bp paired end reads for indel identification. Sequencing depth was generally > 500,000 reads per site. To determine %reads with INDELs, CRISPResso was utilized (http://crispresso.pinellolab.partners.org/) with the following arguments: “CRISPResso -r1 read1.fastq -r2 read2.fastq -a amplicon -g gRNA/off-target site seq -w 40 -q 30 --ignore_substitutions”

## Acknowledgements

We thank Alec R. Nickolls, Thomas Winkler, Sarah Davies and Diego Espinoza for their helpful advice, AbdulAziz AlJanahi for technical assistance, the NHLBI DNA Sequencing and Genomics Core for DNA sequencing, Keyvan Keyvanfar for cell sorting, and Eric Moon for providing AmpliSeq HD sequencing materials. We thank the animal care staff for their support of all animal care and procedures.

This work was supported by the Intramural Program of the National Heart, Lung, and Blood Institute, St. Jude Children’s Research Hospital and ALSAC, National Institutes of Health Common Fund Somatic Cell Genome Editing grant U01HL145793, and the Saudi Arabian Cultural Mission to the United States.

## Author Contributions

Conceptualization, A.A.A., S.Q.T. and C.E.D.; Methodology, K.Y., M.Y.K., D.Y., T.C. and S.Q.T.; Software, A.A.A, S. Cordes and I.T.; Visualization: A.A.A. and L.L.T.; Investigation, A.A.A., C.R.L., S. Chen, T-H.S., I.J., Y.Z., B.L., S.G.H. and J.K.; Resources, Y.L. and B.T.; Writing – Original Draft, A.A.A. and C.E.D.; Writing – Review & Editing, A.A.A., T-H.S, S.G.H, Y.Z., S.Q.T. and C.E.D., Funding Acquisition, C.E.D. and A.A.A.; Supervision, T.C., S.G., S.Q.T. and C.E.D.

## Supplemental Information

**Figure S1.**
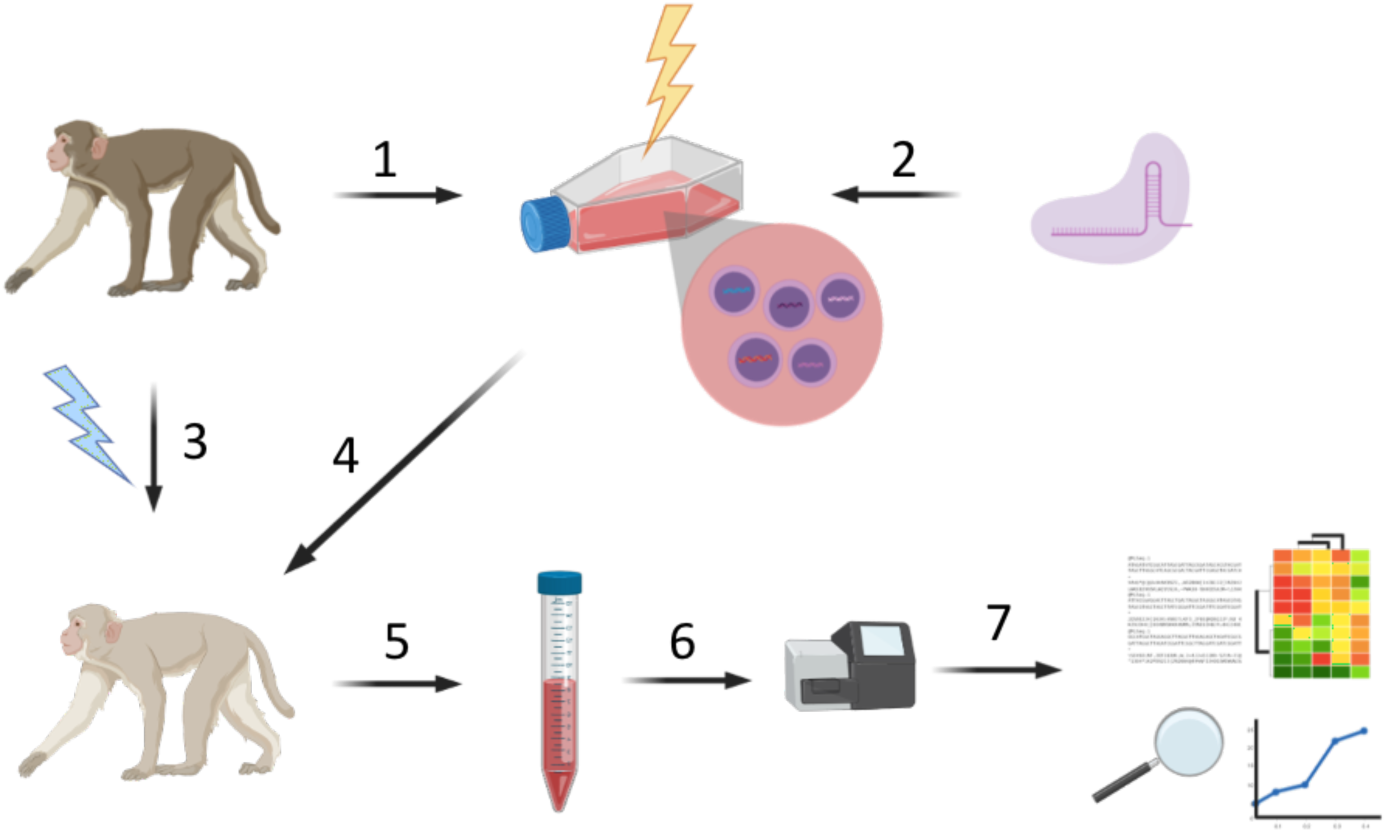
Schematic of autologous HSPC editing in the rhesus macaque model. 1- CD34+ cells are collected from the animal. 2- the collected cells are edited with Cas9+gRNA to create the infusion product. 3- the animal is exposed to radiation to empty the bone marrow niche for the new edited cells. 4- the edited cells are transfused back into the animal. 5- the edited cells start producing cells that populate the peripheral blood, which can be sampled at different time points. 6- DNA from cells collected at different time points can be sequenced in search for edits. 7- sequencing results are analyzed to look for on-target and off-target editing over-time.

**Figure S2.**
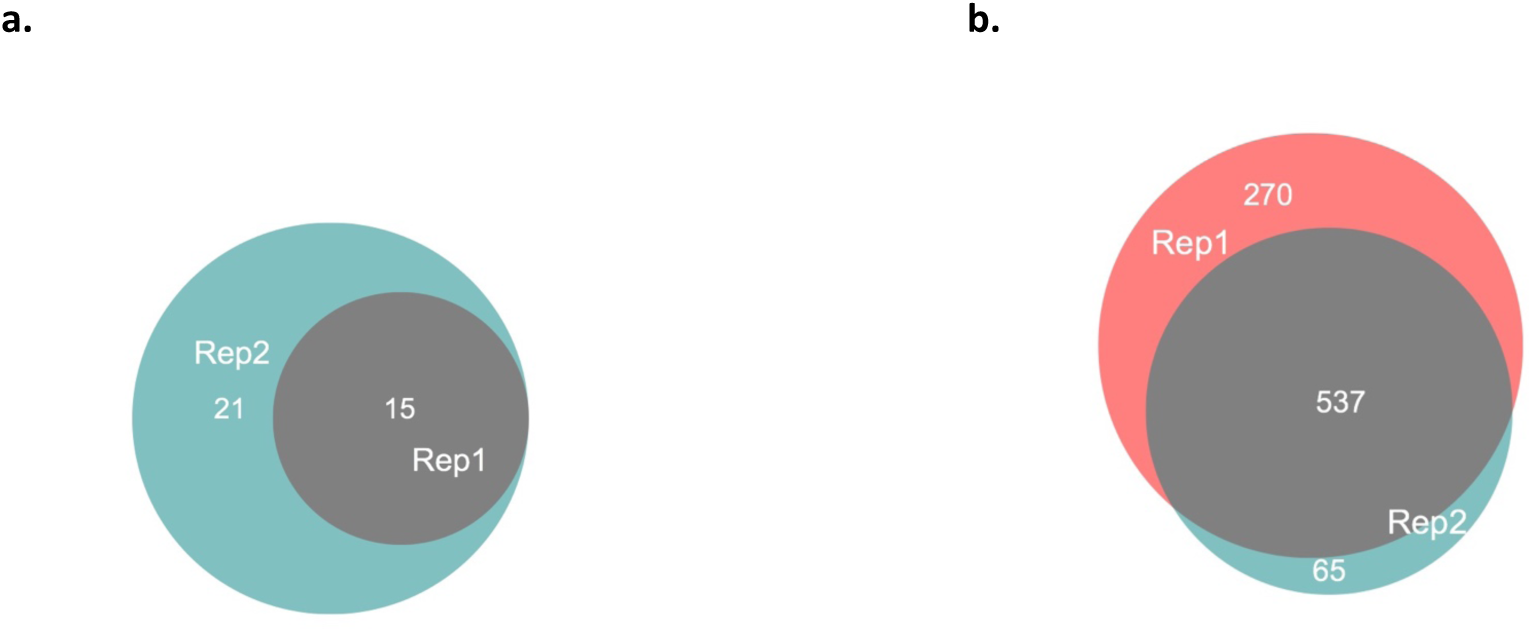
The site overlap between CIRCLE-Seq replicates for TET2 and CD33. **A)** Overlap between the off-target sites predicted by the TET2 CIRCLE-Seq technical replicates performed on animal ZL26. **B)** Overlap between the off-target sites predicted by the CD33 CIRCLE-Seq technical replicates performed on animal ZJ52.

**Figure S3.**
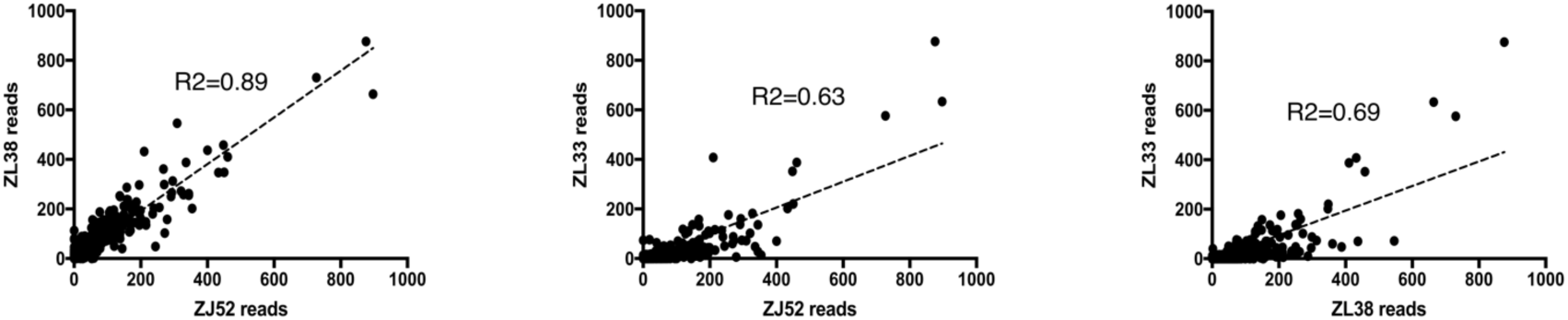
Correlation of read counts for the CIRCLE-Seq detected CD33 off-target sites in 3 animals. Normalized read count plots showing the correlation between the reads for the CD33 off-target sites detected by CIRCLE-seq, 2 animals at a time.

**Figure S4.**
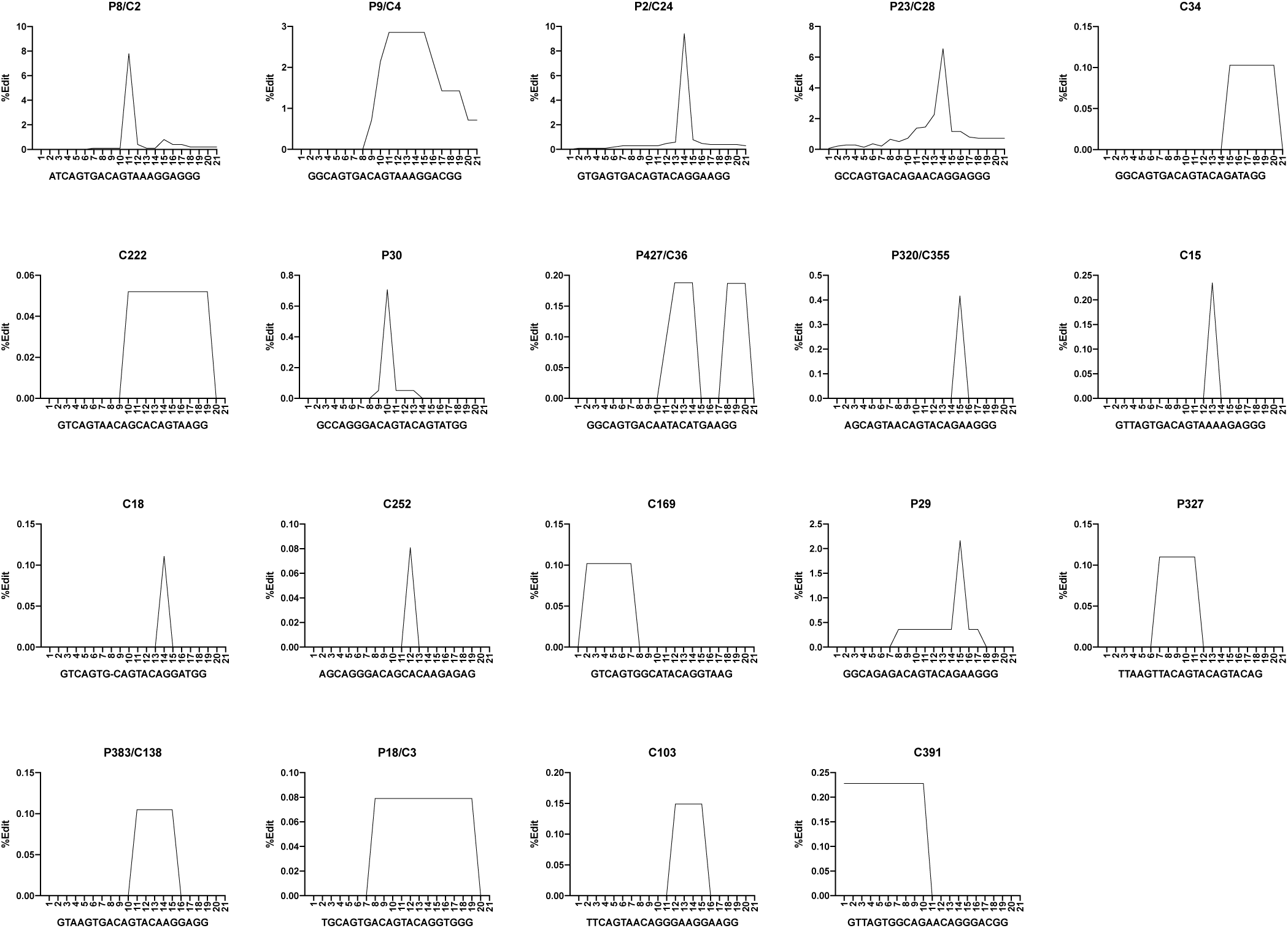
Editing patterns in the 19 *bona fide* CD33 off-target sites. Percent editing in each position of the 21 bps complimentary to the CD33 gRNA. The editing percentages are shown in the highest edited sample for each of the 19 edited off-target sites. All 19 sites showed no detectable variation from the reference in the un-edited samples.

**Table S1.**
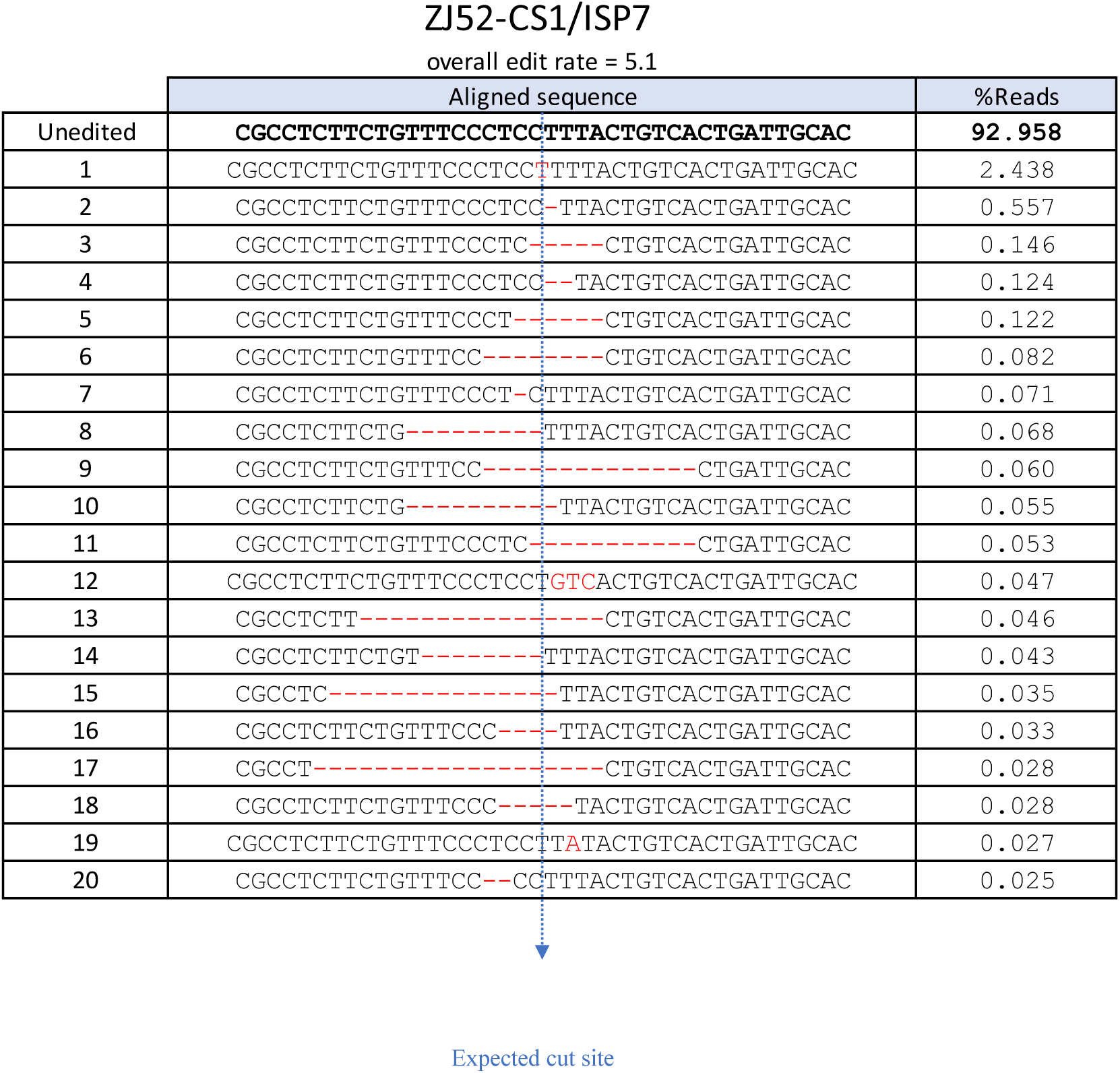
Top 20 edited reads in the infusion product of ZL52 at site ZJ52-CS1/ISP7.

**Table S2.**
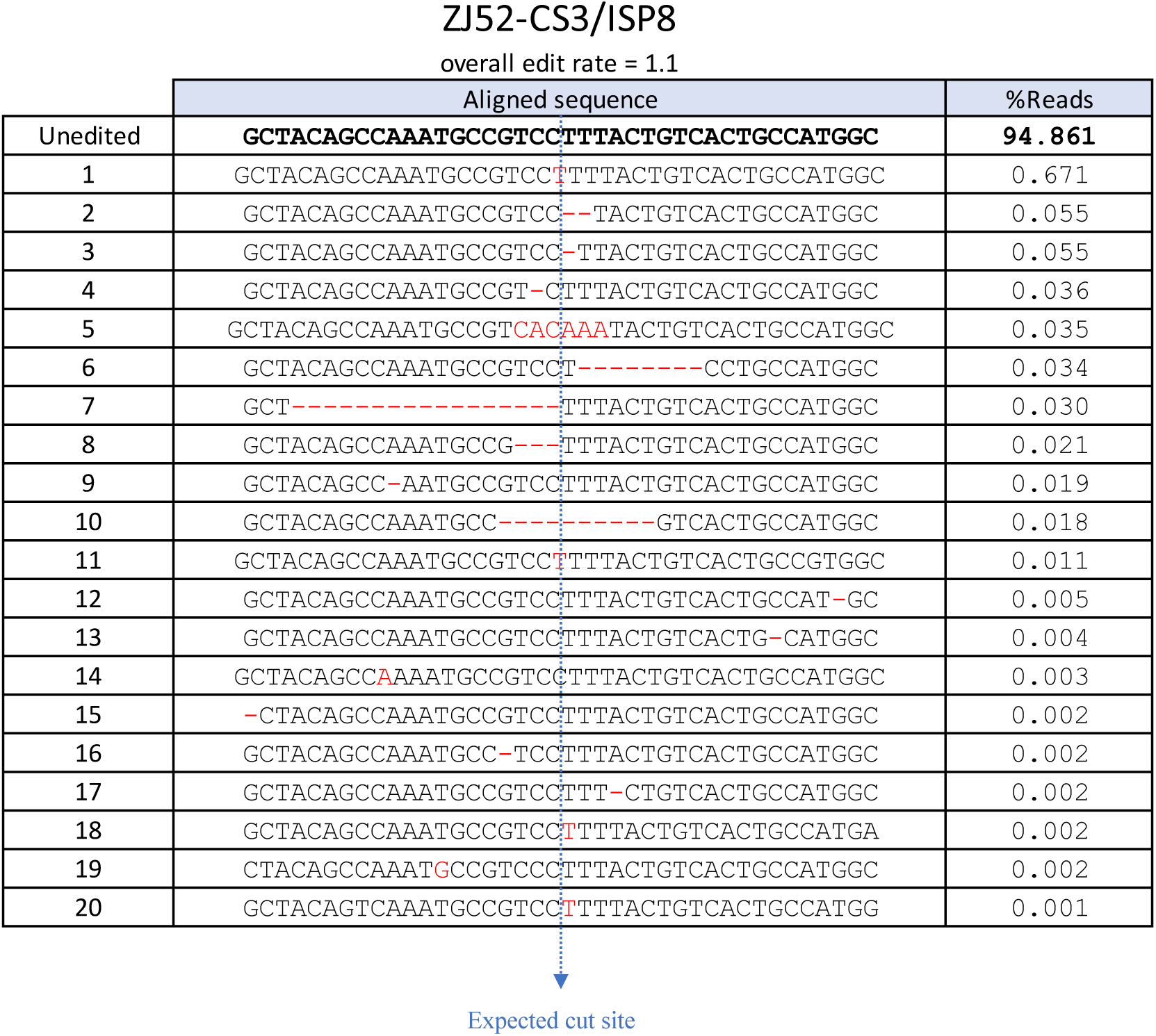
Top 20 edited reads in the infusion product of ZL52 at site ZJ52-CS3/ISP8.

**Table S3.**
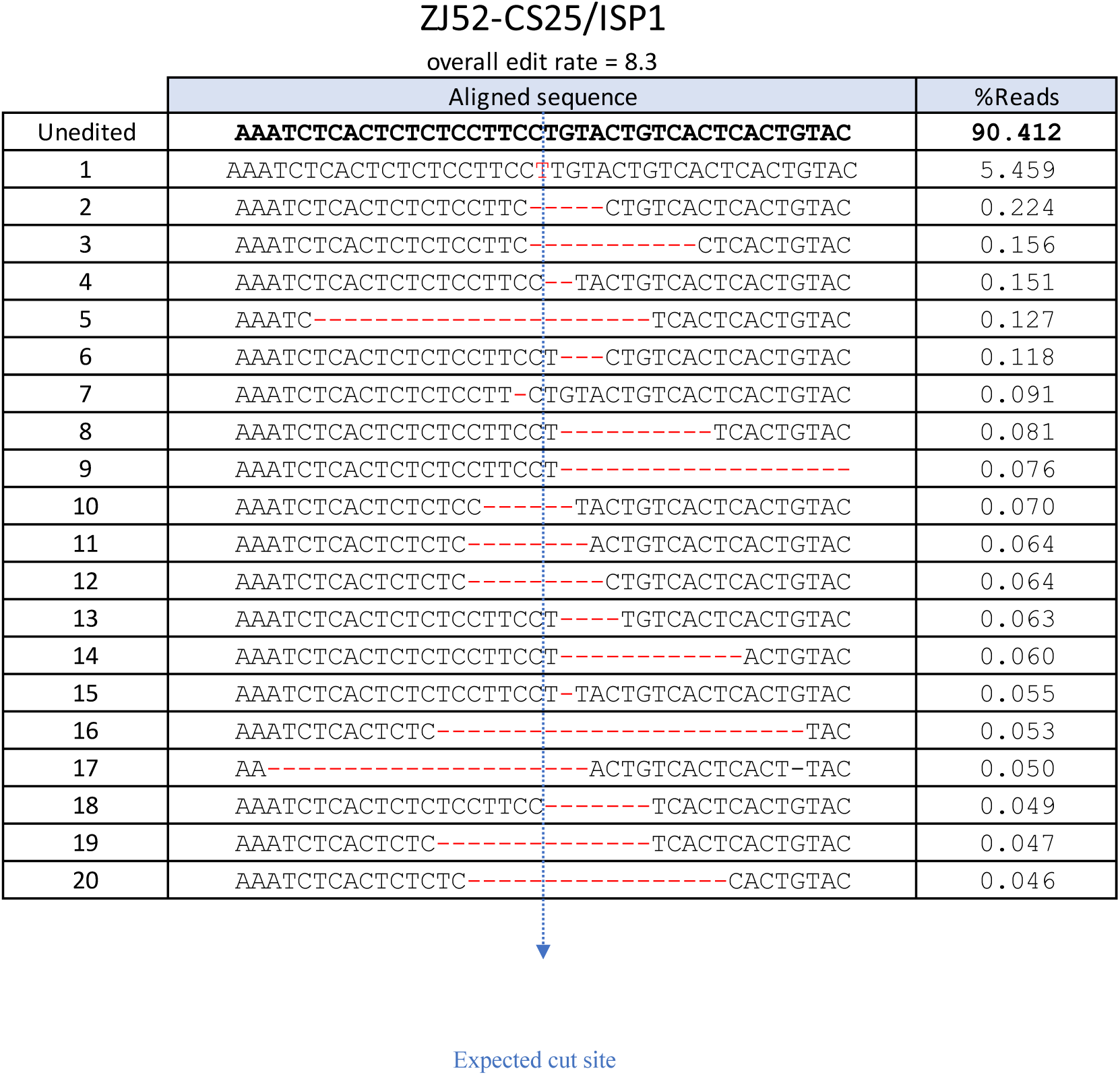
Top 20 edited reads in the infusion product of ZL52 at site ZJ52-CS25/ISP1.

**Table S4.**
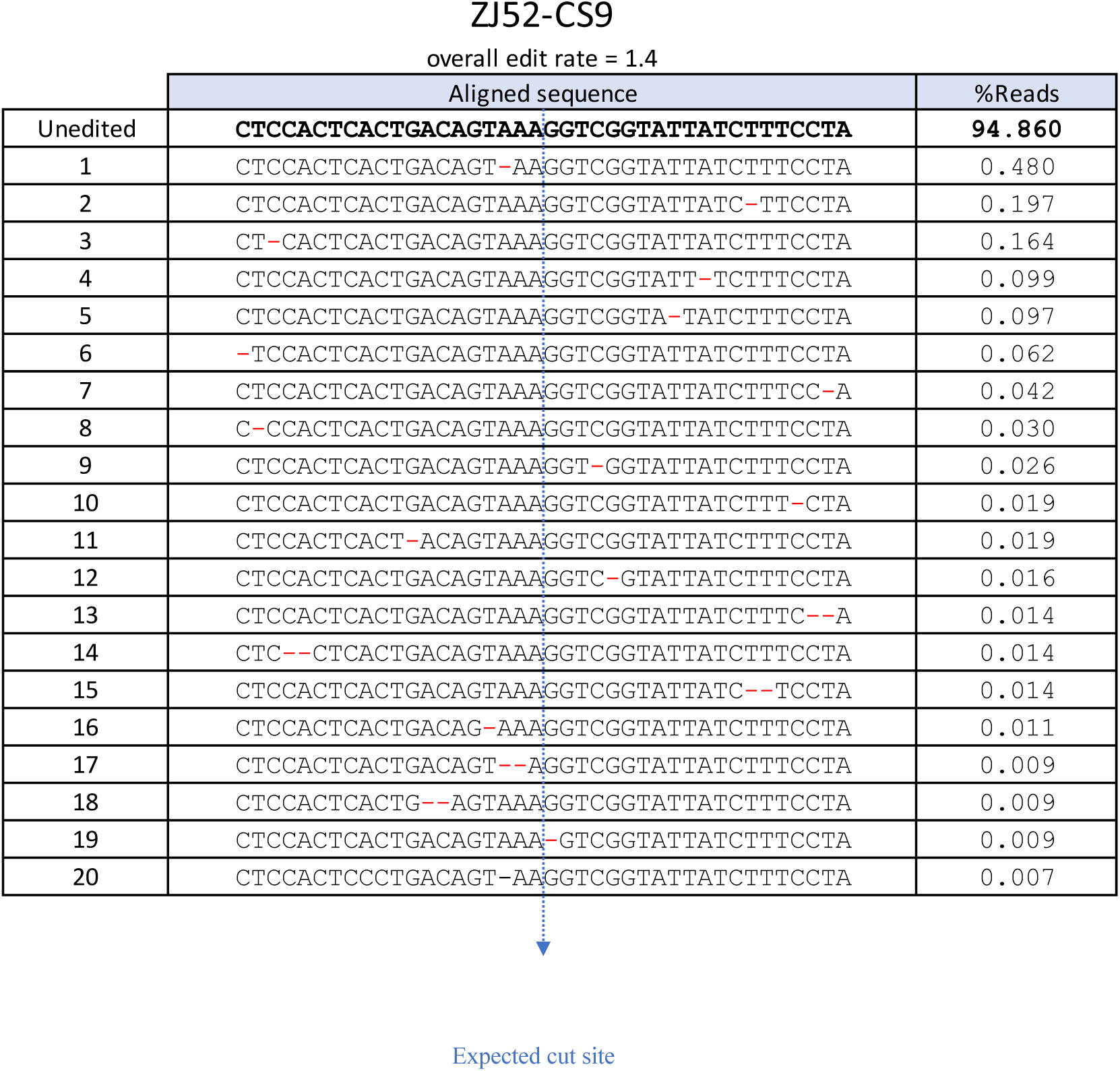
Top 20 edited reads in the infusion product of ZL52 at site ZJ52-CS9.

**Table S5.**
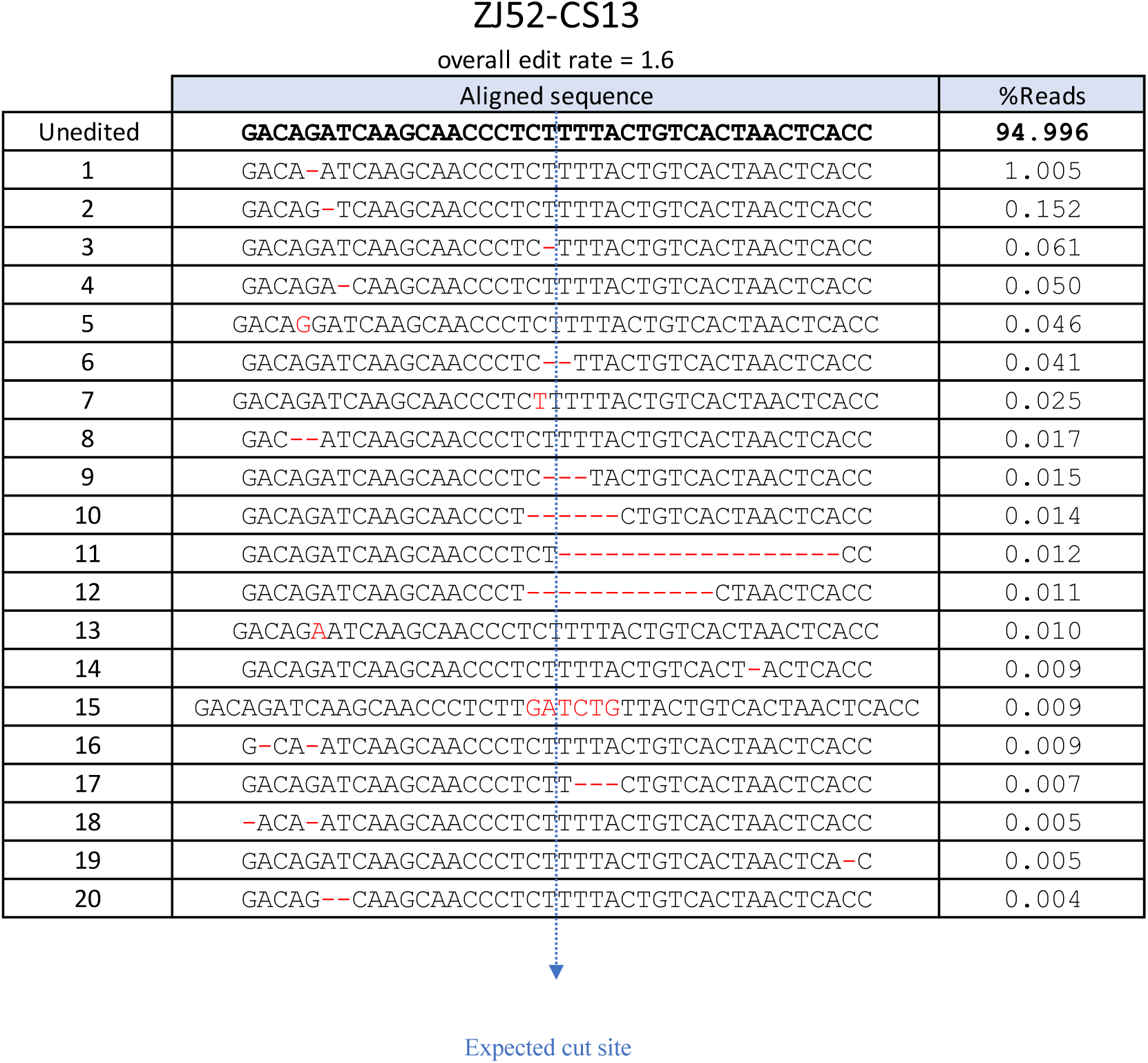
Top 20 edited reads in the infusion product of ZL52 at site ZJ52-CS13.

**Table S6.**
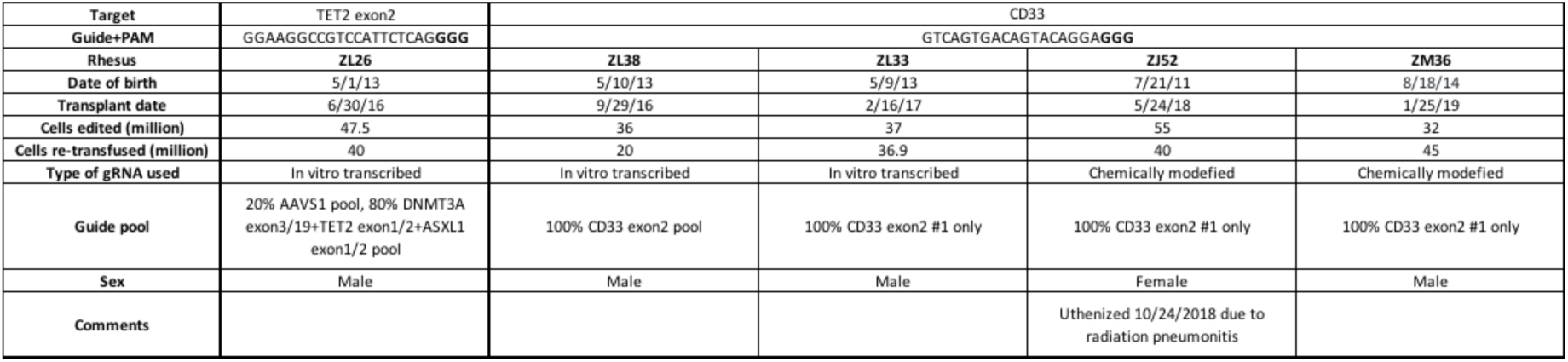
Details of the transplantation and editing parameters for each edited rhesus.

**Table S7.**
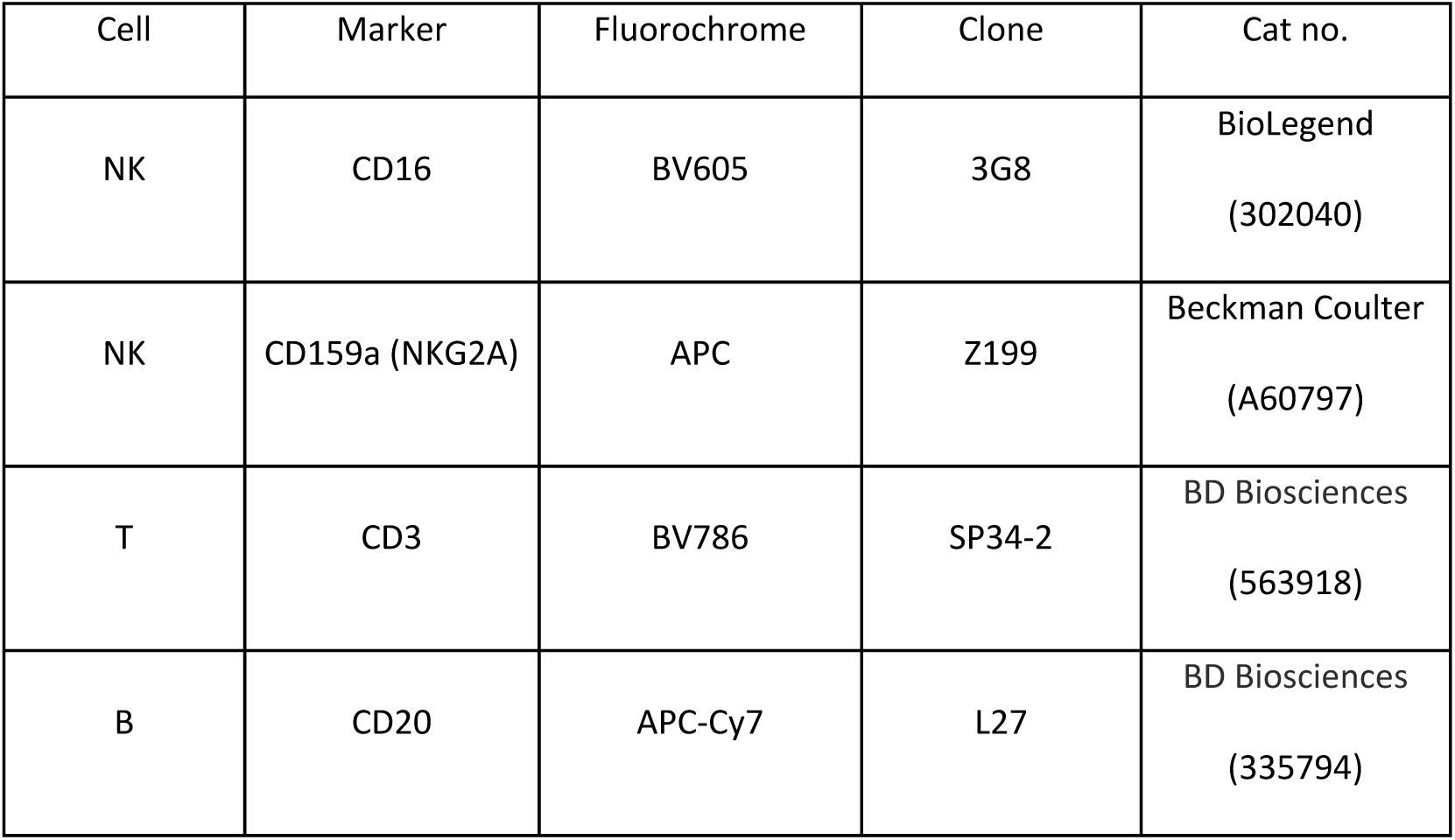
Monoclonal antibody clones used for flow-cytometry sorting.

**Table S8.**
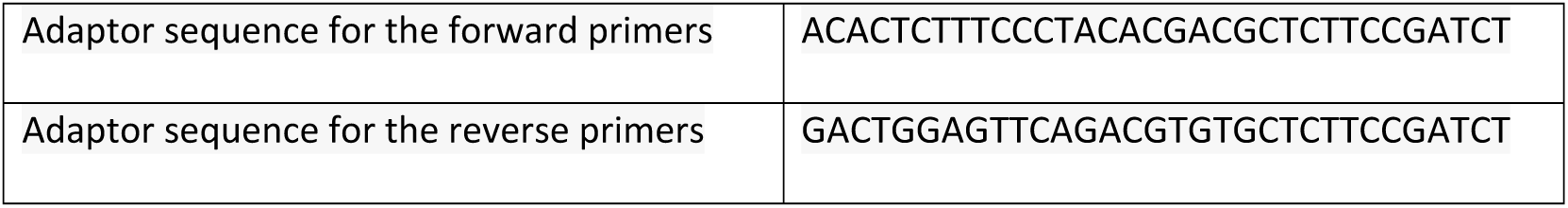
Adaptor sequences for primers of targeted Illumina sequencing library prep.

## References

1. Jinek, M. et al. A Programmable Dual-RNA – Guided. 337, 816–822 (2012).

2. Ran, F. A. et al. Genome engineering using the CRISPR-Cas9 system. Nat. Protoc. 8, 2281–2308 (2013).

3. Akcakaya, P. et al. In vivo CRISPR editing with no detectable genome-wide off-target mutations. Nature 561, 416–419 (2018).

4. Koo, T. & Kim, J.-S. Therapeutic applications of CRISPR RNA-guided genome editing. Brief. Funct. Genomics 16, 38–45 (2017).

5. Kim, M. Y. et al. Genetic Inactivation of CD33 in Hematopoietic Stem Cells to Enable CAR T Cell Immunotherapy for Acute Myeloid Leukemia. Cell (2018) doi:10.1016/j.cell.2018.05.013.

6. Iyer, V. et al. CRISPR off-target analysis in genetically engineered rats and mice. PLoS Genet. 15, 512–514 (2018).

7. Li, C. et al. Trio-Based Deep Sequencing Reveals a Low Incidence of Off-Target Mutations in the Offspring of Genetically Edited Goats. Front. Genet. 9, 1–12 (2018).

8. Lin, Y. et al. CRISPR/Cas9 systems have off-target activity with insertions or deletions between target DNA and guide RNA sequences. Nucleic Acids Res. 42, 7473–7485 (2014).

9. Ayabe, S., Nakashima, K. & Yoshiki, A. Off- and on-target effects of genome editing in mouse embryos. J. Reprod. Dev. 65, 1–5 (2019).

10. Yee, J. Off-target effects of engineered nucleases. FEBS J. 283, 3239–3248 (2016).

11. Martin, F., Sánchez-hernández, S., Gutiérrez-guerrero, A., Pinedo-gomez, J. & Benabdellah, K. Biased and Unbiased Methods for the Detection of Off-Target Cleavage by CRISPR / Cas9: An Overview. Int. J. Mol. Sci. (2016) doi:10.3390/ijms17091507.

12. Haeussler, M. et al. Evaluation of off-target and on-target scoring algorithms and integration into the guide RNA selection tool CRISPOR. Genome Biol. 17, 148 (2016).

13. Tsai, S. Q. et al. GUIDE-seq enables genome-wide profiling of off-target cleavage by CRISPR-Cas nucleases. Nat. Biotechnol. 33, 187–197 (2015).

14. Tsai, S. Q. et al. CIRCLE-seq: a highly sensitive in vitro screen for genome-wide CRISPR–Cas9 nuclease off-targets. Nat. Methods 14, 607–614 (2017).

15. Hsu, P. D. et al. DNA targeting specificity of RNA-guided Cas9 nucleases. Nat. Biotechnol. 31, 827–832 (2013).

16. Hong, S. G. et al. Rhesus iPSC Safe Harbor Gene-Editing Platform for Stable Expression of Transgenes in Differentiated Cells of All Germ Layers. Mol. Ther. 25, 44–53 (2017).

17. Dunbar, C. E. et al. Gene therapy comes of age. Science (80-.). 359, eaan4672 (2018).

18. Larochelle, A. & Dunbar, C. E. Hematopoietic Stem Cell Gene Therapy: Assessing the Relevance of Preclinical Models. Semin. Hematol. 50, 101–130 (2013).

19. Yu, K.-R. et al. A Non-Human Primate CRISPR/Cas9 Model of Clonal Hematopoiesis of Indeterminate Potential Demonstrates Expansion of TET2-Disrupted Clones. (2017).

20. Mali, P. et al. RNA-Guided Human Genome Engineering via Cas9. Science (80-.). 339, 823–826 (2013).

21. Fonslow, B. R. et al. Rational design of highly active sgRNAs for CRISPR-Cas9-mediated gene inactivation. Nat. Biotechnol. 32, 1262–1267 (2014).

22. Fu, Y., Sander, J. D., Reyon, D., Cascio, V. M. & Joung, J. K. Improving CRISPR-Cas nuclease specificity using truncated guide RNAs. Nat. Biotechnol. 32, 279–284 (2014).

23. Shin, T. H. et al. CRISPR/Cas9 PIG-A gene editing in nonhuman primate model demonstrates no intrinsic clonal expansion of PNH HSPCs. Blood 133, 2542–2545 (2019).

24. Ma, X. et al. Analysis of error profiles in deep next-generation sequencing data. Genome Biol. 20, 50 (2019).

25. Akcakaya, P. et al. In vivo CRISPR-Cas gene editing with no detectable genome-wide off-target mutations. bioRxiv (2018) doi: https://doi.org/.

26. Cradick, T. J., Qiu, P., Lee, C. M., Fine, E. J. & Bao, G. COSMID: A Web-based Tool for Identifying and Validating CRISPR/Cas Off-target Sites. Mol. Ther. Nucleic Acids 3, e214 (2014).

27. Buenrostro, J. D., Wu, B., Chang, H. Y. & Greenleaf, W. J. ATAC-seq: A Method for Assaying Chromatin Accessibility Genome-Wide. Curr. Protoc. Mol. Biol. 109, 21.29.1-21.29.9 (2015).

28. Buenrostro, J. D. et al. Integrated Single-Cell Analysis Maps the Continuous Regulatory Landscape of Human Hematopoietic Differentiation. Cell 173, 1535–1548.e16 (2018).

29. Espinoza, D. A. et al. Aberrant Clonal Hematopoiesis following Lentiviral Vector Transduction of HSPCs in a Rhesus Macaque. Mol. Ther. 27, 1074–1086 (2019).

30. Yin, H. et al. Therapeutic genome editing by combined viral and non-viral delivery of CRISPR system components in vivo. Nat. Biotechnol. 34, 328–333 (2016).

31. Ding, Q. et al. Permanent Alteration of PCSK9 With In Vivo CRISPR-Cas9 Genome Editing. Circ. Res. 115, 488–492 (2014).

32. Nelson, C. E. et al. In vivo genome editing improves muscle function in a mouse model of Duchenne muscular dystrophy. Science (80-.). 351, 403–407 (2016).

33. Long, C. et al. Postnatal genome editing partially restores dystrophin expression in a mouse model of muscular dystrophy. Science (80-.). 351, 400–403 (2016).

34. Wu, Y. et al. Correction of a Genetic Disease in Mouse via Use of CRISPR-Cas9. Cell Stem Cell 13, 659–662 (2013).

35. Amoasii, L. et al. Gene editing restores dystrophin expression in a canine model of Duchenne muscular dystrophy. Science (80-.). 362, 86–91 (2018).

36. Veres, A. et al. Low Incidence of Off-Target Mutations in Individual CRISPR-Cas9 and TALEN Targeted Human Stem Cell Clones Detected by Whole-Genome Sequencing. Cell Stem Cell 15, 27–30 (2014).

37. Chen, Y. et al. Functional disruption of the dystrophin gene in rhesus monkey using CRISPR/Cas9. Hum. Mol. Genet. 24, 3764–3774 (2015).

38. Sui, T. et al. A novel rabbit model of Duchenne muscular dystrophy generated by CRISPR/Cas9. DMM Dis. Model. Mech. 11, (2018).

39. Wang, S. et al. No off-target mutations in functional genome regions of a CRISPR/Cas9-generated monkey model of muscular dystrophy. J. Biol. Chem. 293, 11654–11658 (2018).

40. Iyer, V. et al. No unexpected CRISPR-Cas9 off-target activity revealed by trio sequencing of gene-edited mice. PLOS Genet. 14, e1007503 (2018).

41. Anderson, K. R. et al. CRISPR off-target analysis in genetically engineered rats and mice. Nat. Methods 15, 512–514 (2018).

42. Aryal, N. K., Wasylishen, A. R. & Lozano, G. CRISPR/Cas9 can mediate high-efficiency off-target mutations in mice in vivo. Cell Death Dis. 9, 1099 (2018).

43. Komor, A. C., Badran, A. H. & Liu, D. R. CRISPR-Based Technologies for the Manipulation of Eukaryotic Genomes. Cell 169, 559 (2017).

44. Tsai, S. Q., Joung, J. K., Capecchi, M. & Evans, M. Defining and improving the genome-wide specificities of CRISPR –Cas9 nucleases. Nat. Publ. Gr. 17, 300–312 (2016).

45. Kuscu, C., Arslan, S., Singh, R., Thorpe, J. & Adli, M. Genome-wide analysis reveals characteristics of off-target sites bound by the Cas9 endonuclease. Nat. Biotechnol. 32, 677–683 (2014).

46. Fu, Y. et al. High-frequency off-target mutagenesis induced by CRISPR-Cas nucleases in human cells. Nat. Biotechnol. 31, 822–826 (2013).

47. Kim, S., Kim, D., Cho, S., Kim, J. & Kim, J.-S. Highly Efficient RNA-guide genome editing… Genome Res. 24, 1012–1019 (2014).

48. Gao, X. et al. Treatment of autosomal dominant hearing loss by in vivo delivery of genome editing agents. Nature 553, 217–221 (2018).

49. Tsai, S. Q. et al. GUIDE-seq enables genome-wide profiling of off-target cleavage by CRISPR-Cas nucleases. Nat Biotechnol 33, 187–197 (2015).

50. Yang, L. et al. Targeted and genome-wide sequencing reveal single nucleotide variations impacting specificity of Cas9 in human stem cells. Nat. Commun. 5, 5507 (2014).

51. McInerney, P., Adams, P. & Hadi, M. Z. Error Rate Comparison during Polymerase Chain Reaction by DNA Polymerase. Mol. Biol. Int. 2014, 1–8 (2014).

52. Filges, S., Yamada, E., Ståhlberg, A. & Godfrey, T. E. Impact of Polymerase Fidelity on Background Error Rates in Next-Generation Sequencing with Unique Molecular Identifiers/Barcodes. Sci. Rep. 9, 1–7 (2019).

53. de Paz, A. M. et al. High-resolution mapping of DNA polymerase fidelity using nucleotide imbalances and next-generation sequencing. Nucleic Acids Res. 46, e78 (2018).

54. Ihry, R. J. et al. p53 inhibits CRISPR–Cas9 engineering in human pluripotent stem cells. Nat. Med. 24, 939–946 (2018).

55. Donahue, R. E., Kuramoto, K. & Dunbar, C. E. Large Animal Models for Stem and Progenitor Cell Analysis. Curr. Protoc. Immunol. 69, 1–29 (2005).

